# Reconstructing bat antiviral immunity using epithelial organoids

**DOI:** 10.1101/2024.04.05.588241

**Authors:** Max J. Kellner, Patrick Zelger, Vanessa Monteil, Gang Pei, Masahiro Onji, Komal Nayak, Matthias Zilbauer, Anne Balkema-Buschmann, Anca Dorhoi, Ali Mirazimi, Josef M. Penninger

**Author notes:** Correspondence (M.J.K), (J.M.P).

## Abstract

Bats are remarkably resilient to viruses with pandemic potential. To resolve largely unknown molecular mechanisms governing their exceptional antiviral immunity, we established an organoid platform to model the entire respiratory airway and intestinal epithelium of the important viral reservoir species *Rousettus aegyptiacus* (Egyptian fruit bat). These bat organoids exhibit an unexpected diversity of cell types and support replication of highly pathogenic zoonotic viruses including Marburg virus (MARV) and MERS-Coronavirus. Following virus infection, bat organoids unleash a strong interferon response, uniquely regulated through virus-dependent and virus-independent mechanisms. By contrast, MARV infected human organoids fail to induce an antiviral gene response and express pro-inflammatory cytokines after interferon stimulation, revealing important molecular differences between bats and humans with implications for lethal Marburg virus infections in primates. These data provide the most comprehensive organoid platform in bats to decode species-specific differences and uncover fundamental principles of bat disease resilience to emerging viruses with pandemic potential.

## Introduction

Bats comprise the second most diverse order in the mammalian kingdom and have evolved a set of unique physiological features since their divergence from a common ancestor more than 50 million years ago^1^. As the sole mammal capable of self-powered flight, multiple bat species are long-lived with little to no sign of age-related tumorigenesis^2–5^. The significance of bats to human health is underscored by their unique ability to host and tolerate pathogens that are highly virulent to humans and non-human primates^6–8^. For example, bats have been identified or suspected to be the natural reservoir for viruses of pandemic concern including Betacoronaviruses related to MERS^9, 10^, SARS^11^ and SARS-CoV-2^12^, hemorrhagic fever Filoviruses (Ebola virus^13^ and Marburg virus^14^), or highly pathogenic Paramyxoviruses (Hendra virus^15^ or Nipah virus^16, 17^). Understanding the molecular principles that govern the biology of bats has motivated on-going research efforts to perform genomic analyses in bats and establish experimental model systems^18–21^, sometimes resulting in contradictory findings: Expansion, contraction, or constitutive expression of genes from the interferon system, absence of killer cell immunoglobulin-like receptors (KIRs) in some bat genomes, dampened inflammation or genetic changes including loss of key signaling proteins (STING1, NLRP1, CARD8)^22–26^. Despite these recent insights into the molecular responses of bats to virus infection, much remains unknown about how bats establish resilience against infectious diseases of human pandemic potential.

Experimental research in bats remains challenging, owing to the limited access to bat breeding colonies, their protection status, as well as a general lack of commercial reagents tested and optimized for bat species^27^. Almost all functional genetic studies in bats have been performed in only a limited number of immortalized cell lines from three different species (PaKiT^21^ (*Pteropus Alecto*), RoNi/7^20^ (*Rousettus aegyptiacus*), R06E^28^ (*Rousettus aegyptiacus*), and Efk3^29^(*Eptesicus fuscus*)). While bat cell lines provide suitable means to study fundamental functions *in vitro*, they often do not reflect the cell-type complexity and associated gene expression patterns found in tissues, including the expression of virus entry factors. Moreover, immortalized cell lines may exhibit variability in their response to stimuli^30^. It is therefore imperative to develop physiologically relevant *in vitro* models to study the complex biology of bats.

Airway and gastrointestinal epithelia are major entry sites for RNA viruses in mammals. These mucosal surfaces constitute the first line of antiviral defense and are an integral part of the innate immune system^31^. In humans, stem cell-derived organoids have been used extensively to assess key aspects of virus infections, host cell responses, or to test drugs and vaccines^32^. A small number of bat organoids has been established so far, including intestinal organoids from *Rhinolophus sinicus*^33^, *Artibeus jamaicensis*^34^ and *Rousettus leschenaultii*^35^, as well as tracheal organoids from *Eonycteris spelaea*^36^ and *Carollia perspicillata*^37^. However, these pioneering studies have critically lacked any in depth molecular characterization and functional genetic perturbations of naïve or infected organoids, likely due to limitation in fully annotated reference genomes. Hence, these models are still largely inaccessible to a wider research community. We therefore set out to generate tools and molecular resources to democratize access to bat research, critically needed considering frequent bat to human spillover of viruses of pandemic potential and the potential of bats, as the second most divers and evolutionary successful mammalian order, to uncover novel biology even beyond infectious biology.

Here we present a sustainable organoid platform to model distinct respiratory tract airway and small intestinal epithelia of *Rousettus aegyptiacus* (Egyptian fruit bat), a natural reservoir for multiple virus species, such as the highly lethal Marburg virus (MARV)^14^. Through deep characterization of organoids, including single-cell RNA-sequencing, and infection with multiple emerging zoonotic RNA viruses, we provide novel insights into the cell-type diversity and innate antiviral immunity of the *Rousettus aegyptiacus* mucosal epithelium. In particular, we show that bat organoids are able inititate an antiviral response following MERS-CoV and MARV infection, and identified an amplifying circuit of the type-III interferon response in bats. Our data uncover fundamental principles of bat innate resilience to virus infections that contrast to humans and could be utilized to translate protective bat immunity to human health.

## Results

### A sustainable bat organoid platform

To advance bat research and democratize access to complex bat *in vitro* models, we established a robust protocol for the generation of adult-stem cell derived organoids from airway and small intestinal tissue of *Rousettus aegyptiacus.* The Egyptian fruit bat is a frugivorous bat of the family of *Pteropodidae* mainly found in Africa that can harbor multiple virus families, including human pathogens: diverse strains of Marburg virus (MARV)^14^, Sosuga virus (SOSV)^38^, Kasokero virus (KASV)^39^ or bat H9N2 Influenza-A virus^40^ (Figure 1A, B). One major bottleneck we encountered was access to bat tissue. Therefore, following multiple pilot trials we established a protocol for effective cryopreservation (see methods) allowing us to maintain primary bat tissues that can be readily shipped to other sites and subsequently used as starting material for our organoid protocols (Figure 1C). This tissue cryopreservation should be also feasible for wild-caught bats, following all protection guidelines and Nagoya protocols. We also established a cryopreservation protocol for already expanded bat organoids as an additional source for distribution and access (Figure 1C). These novel protocols and organoid platforms should readily provide access to complex bat *in vitro* models to a wider research community.

**Figure 1.**
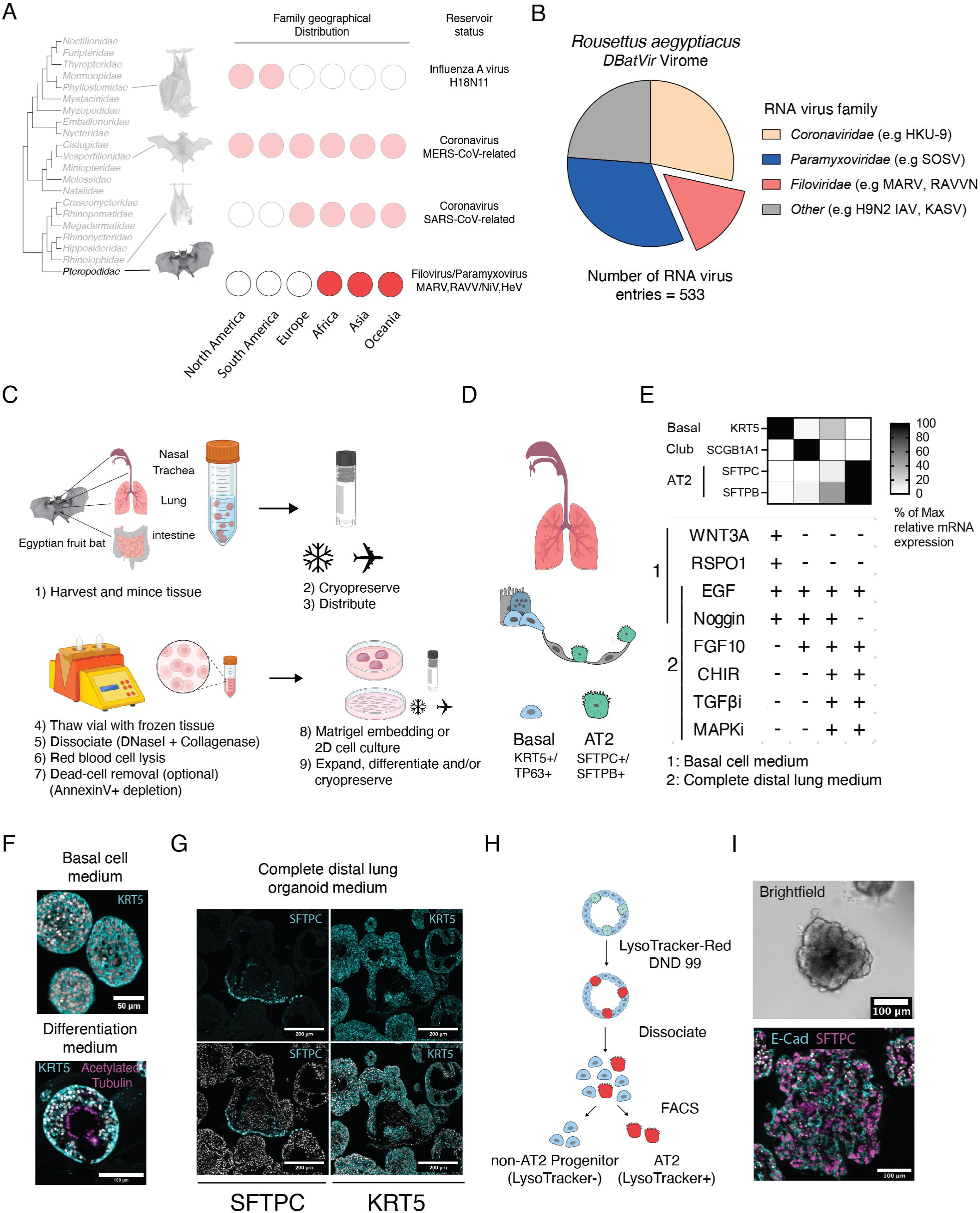
Self-replicating organoids from Egyptian fruit bats. A) Phylogenetic tree of *Chiroptera* families, geographic distribution and reservoir status. Information content regarding bat dispersion adapted from^108^ B) RNA virus species identified in *Rousettus aegyptiacus*. Data was derived from the DBatVir^109^ database and represent individual entries grouped by virus family. C) Workflow for generating bat organoids from cryogenically preserved tissue. D) Schematic of airway progenitor cells of the airway epithelium *in vivo*. E) RT-qPCR normalized expression values for specific marker genes (basal, club or AT2 cells) are shown for organoids grown in basal medium with addition of depicted factors. Expression was normalized to the maximum relative mRNA expression (2^-dCT) value found for each gene and averaged across replicates. F) Top: Immunofluorescence anti-KRT5 staining of expanding basal cell organoids. Bottom: Immunofluorescence anti-KRT5 and anti-acetylated tubulin staining of basal cell organoids differentiated in 3D. G) Immunofluorescence images of distal lung organoids cultured in *complete distal lung expansion medium* prior to FACS enrichment of LysoTracker-Red DND-99 dye positive cells. Anti-SFTPC or Anti-KRT5 antibodies were used to stain AT2 and basal cells in the same region of organoid sections (top) and counterstained with DAPI (bottom). H) Schematic of FACS AT2-cell enrichment using LysoTracker-Red DND-99. I) Brightfield (top) and immunofluorescence (bottom) image of alveolar organoids post LysoTracker-dye FACS enrichment stained with anti-SFTPC and anti-E-cadherin antibody.

### Reconstructing the entire respiratory epithelial tree of Egyptian fruit bats *in vitro*

We hypothesized that Egyptian fruit bat adult epithelial stem cells require similar growth factors for maintenance as in other mammals^41^. The airway is comprised of at least two distinct self-replicating adult stem-cell progenitor cell types, basal stem cells expressing cytokeratin-5 (KRT5) and p63 (TP63) and surfactant protein-C/B expressing (SFTPC, SFTPB) alveolar type II stem cells (AT2)^42^. While basal stem cells are mainly found in the nasal respiratory epithelium and conducting airway of trachea and lung, AT2 progenitors are located in the distal alveolar epithelium (Figure 1D). We therefore determined factors required to maintain these airway progenitor cells *in vitro*^43–45^. KRT5+ basal stem cells from *Rousettus aegyptiacus* nasal, tracheal, and distal lung epithelium could be maintained in media containing growth factors that activate EGF and FGF receptor signaling (EGF, FGF10) and induce the WNT pathway (using R-spondin1), while blocking BMP receptor signaling (Noggin) (Figure 1E). Of note, while WNT3A was not required for long-term maintenance, inhibition of TGF-β/Smad signaling using A83-01 improved long-term airway basal cell organoid growth. Blockage of EGFR and/or FGFR signaling through small molecules (PD153035 for EGFR, Futibatinib for FGFR), inhibited organoid growth (Supplemental Figure S1A). Expanding nasal, tracheal, and distal lung KRT5+ basal cell organoids displayed a compact morphology and upon differentiation formed a lumen with inwards facing beating cilia (Figure 1F, Supplemental Figure S1B, Supplemental movie 1). These structures are highly reminiscent of human nasal and large airway organoids as well as bat tracheal organoids established previously^36, 37, 44, 46^. Basal cell airway organoids from the upper (nose) and lower (trachea, conducting airways) respiratory tract could be expanded for more than 6 months and retained prototypic transcription factor expression *in vitro*, such as SIX3 for the upper and IRX2 for the lower airway^47^ (Supplemental Figure S1C).

Distal lung organoids containing KRT5+ basal cells and SFTPC+ AT2 cells required EGF, FGF10 and small molecules for activating WNT (CHIR9902), as well as inhibition of p38 MAPK (BIRB-796) and the TGF-β type I receptor signaling (SB-431542) (Figure 1E, G, Supplemental Figure S1D). Interestingly, we found that Noggin provides a key factor for growth and maintainance of distal lung organoids containing KRT5+ basal stem cells and SFTPC+ AT2 cells. SFTPC expression was gradually lost during passaging, likely due to out-competition of SFTPC+ AT2 alveolar stem cells by KRT5+ basal stem cells (Supplemental Figure S1E). To overcome the loss of SFTPC+ alveolar stem cells, we employed a strategy for FACS-based separation of basal and alveolar stem cells through staining of alveolar type II pneumocytes (AT2) using LysoTracker-Red (Figure 1H, Supplemental Figures S1F, G, H). LysoTracker-Red is a non-toxic fluorescence dye that stains the lysosomal-related lamellar bodies specific to AT2 cells^48^. This procedure allowed for the enrichment of AT2 cells and the establishment of bat alveolar organoids that contained self-replicating SFTPC+ alveolar AT2 cells (Figure 1I). Thus, we have generated a robust pipeline for growth and differentiation of nasal, tracheal, distal lung and alveolar organoids from frozen tissue, thereby reconstructing the entire respiratory airway tree of the Egyptian fruit bat *in vitro*.

### Single-cell RNA profiling of Egyptian fruit bat airway organoids

The contemporary view of cell types found in the airway epithelium include basal stem cells, which differentiate in secretory goblet or multi-ciliated cells via suprabasal or club cells^42^. Basal cells can further differentiate into rare cell populations such as ionocytes, endocrine and brush cells. Specialized pulmonary epithelial cells of the distal lung include AT2 cells, which self-replicate or differentiate into AT1 pneumocytes^42^ (Figure 2A). To characterize the cell type diversity of bat airway organoids, we first differentiated nasal and distal lung basal-stem cell derived 3D-cultures at the air liquid interface (ALI) to obtain organotypic cultures with apical-polarization (Acetylated tubulin+ ciliated cells) basal (KRT5+ basal stem cells) and beating cilia facing the air surface (Figure 2B, Supplemental Figure S2A, Supplemental movie 2). The cultures where then dissociated alongside Lysotracker-dye enriched alveolar organoids, and subsequently subjected to 10x droplet-based single-cell RNA-sequencing, resulting in ∼40.000-sequenced cells after quality control and filtering (Supplemental Figure 2A). In parallel, we differentiated human nasal and bronchial epithelial cells at the air-liquid interface under identical conditions and prepared single-cell RNA-sequencing libraries for side-by-side comparisons of differentiated respiratory human and bat cultures.

**Figure 2.**
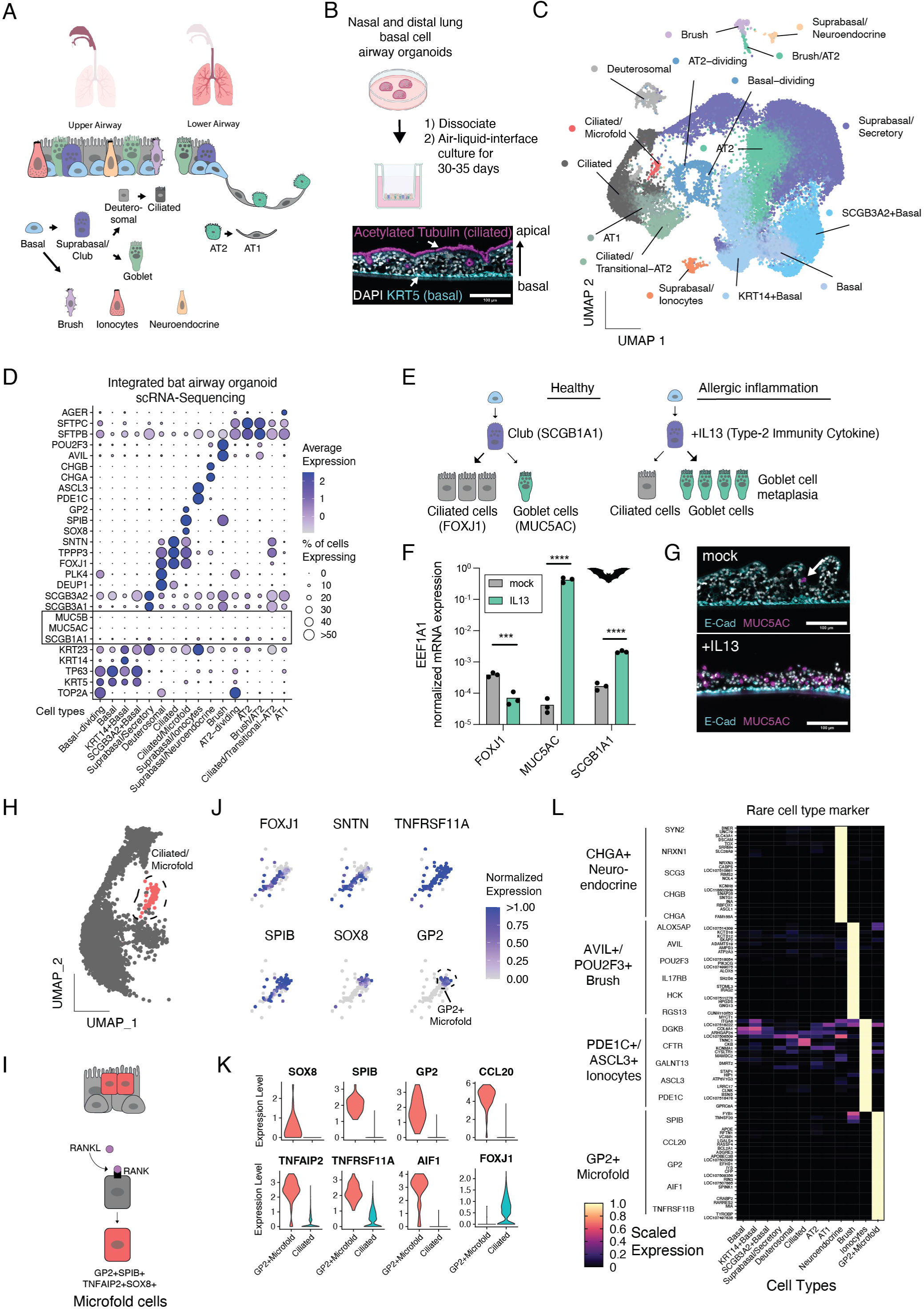
Single-cell profiling of bat airway organoids. A) Schematic depicting major cell types found in the upper and lower airway epithelium of mammals. B) Top: Schematic illustrating workflow to generate air-liquid interface (ALI) cultures from basal cell organoids. Bottom: Immunofluorescence image of ALI cultures showing basal stem cells (anti-KRT5) or apical ciliated cells (anti-acetylated tubulin) and nuclei stained with DAPI. C) UMAP plot of integrated scRNA-sequencing dataset showing cell type clusters of differentiated bat basal cell and alveolar airway organoids. D) DotPlot analysis showing the average expression of markers for each cell cluster. The dot size shows the percentage of an individual cell type expressing a given marker while the color intensity showing the average expression value. Dot sizes were set to a maximum percentage of cells expressing the feature (dot size) of 50%. Genes expressed by more than 50% of cells have the same dot size. E) Schematic showing effect of IL-13 on airway cell differentiation. F) RT-qPCR analysis of reference gene normalized expression of FOXJ1, MUC5AC or SCGB1A1 in bat nasal ALI cultures treated with 10 ng/ml recombinant human IL-13, initiated 10 days after ALI differentiation for 15 days. Each dot represents an individual sample. G) Immunofluorescence anti-MUC5AC and anti-E-cadherin staining of sections from bat nasal ALI cultures treated with 10 ng/ml recombinant human IL-13, initiated 10 days after ALI differentiation for 15 days. Untreated ALI are shown as control. Arrow indicates a single MUC5AC positive cell in control organoids. H) UMAP plot for ciliated and mixed ciliated/microfold cell clusters. The mixed cluster is highlighted in red. I) A schematic for microfold cell formation involving RANK/RANKL. J) FeaturePlot showing the expression of ciliated (FOXJ1, SNTN) and microfold marker (TNFRSF11A, SPIB, SOX8, GP2) in each cell for the mixed ciliated/microfold cell cluster in the integrated UMAP space. The color indicates the normalized expression level with a maximum expression color cutoff set to 1. GP2+ cells are highlighted. K) VlnPlot analysis of microfold marker (SOX8, SPIB, GP2, TNFAPI2, TNFRSF11A, CCL20, TNFAPI2, AIF1) and FOXJ1 shown for GP2+ microfold cells and ciliated cells. The expression distribution is derived from individual cells. L) Heatmap of differentially expressed genes for each rare cell type population in comparison to a collection of other remaining cell types found in bat airway organoids. Expression counts for each gene of each cell type were normalized to the maximum observed value. Previously confirmed specific marker genes for each cell type are highlighted.

To annotate cell types from the upper and lower airway, we performed integrative single cell sequencing analyses using SeuratCCA^49–51^ (Supplemental Table 1). Cells from bat organoids clustered by cell-type rather than organoid model or individuals (Figure 2C, Supplemental Figure S2B, C). We also identified two distinct basal cell populations characterized by high KRT5/TP63/KRT14 expression, compared to KRT5-low/TP63-high/SCGB3A2+ basal cells found in the distal lung (Supplemental Figure S2D). These findings are in line with recently published single-cell transcriptomic data from human distal lung tissue^52^. Suprabasal cells were identified by decreased expression of the basal stem cell markers (KRT5, TP63) and high expression of KRT23 or SCGB3A1/A2 in the upper and lower airway, respectively (Figure 2D, Supplemental Figure S2D). In nasal and distal lung air-liquid-interface (ALI) cultures, we further identified secretory cells (SCGB3A1 or SCGB3A2), deuterosomal cells (TOP2A, PLK4, DEUP1, FOXJ1), and ciliated cells (FOXJ1, TPPP3, SNTN). In Lysotracker-dye positive alveolar organoids, we confirmed the expected enrichment of SFTPC+SFTPB+ AT2 as well as AGER+ HOPX+ CAV1 AT1 cells, while KRT5+/TP63+ basal stem cells were largely depleted (Supplemental Figure S2D, E).

Surprisingly, we observed little to no expression of club/clara cells (SCGB1A1) or goblet cells (MUC5AC/MUC5B) in bat nasal or lung organoids cultured at the air-liquid interface (Figure 2D, Supplemental Figure S2D). This is in stark contrast to human nasal and bronchial basal cell derived air-liquid interface organoids, prepared in parallel using identical culture conditions (Supplemental Figure S3A, B). Upon examining potential molecular differences explaining this discrepancy, we found low to absent levels of SPDEF in our bat air liquid interface organoids (Supplemental Figure S3C). SPDEF is the master regulator of secretory cell fate^53^. In the human nasal and bronchial epithelial air-liquid interface organoid single-cell RNA-sequencing dataset, SPDEF is highly expressed, including in basal stem cells and muco-ciliary precursor cells (Supplemental Figure S3D). To test whether Egyptian fruit bat nasal basal stem cells have the capacity to differentiate into secretory goblet cells *in vitro*, naturally occurring in human airway organoids, we stimulated bat nasal air-liquid interface cultures with the type-2 immunity associated cytokine IL-13. In humans, IL-13 exposure drives goblet cell metaplasia resulting in an altered ciliated-to-secretory-cell ratio^54^ (Figure 2E). IL-13 treatment of bat nasal air-liquid interface cultures triggered significantly increased expression of SCGB1A1 (secretory cell marker) mRNA and a more than 1000-fold increase in the goblet cell marker MUC5AC, whereas expression of the ciliated cell marker FOXJ1 was decreased (Figure 2F). Immunofluorescence staining of IL-13 treated bat nasal air-liquid interface cultures confirmed an abundance of MUC5AC+ goblet cells as compared to the mock treated condition (Figure 2G). This data demonstrate that goblet cells are scarce in respiratory epithelial air liquid interface cultures of Egyptian fruit bats, yet basal cells retain their ability to differentiate into MUC5AC+ secretory goblet cells.

We also found rare AVIL+POU2F3+ brush cells in the bat nasal and lung air liquid interface organoid cultures (Figure 2D; Supplemental Figure 3E). These cells are important mediators of innate immunity through sensing of pathogens and produce cytokines such as IL-25 and cysteinyl leukotrienes (CysLTs)^55^. We further identified cells expressing markers of ionocytes (ASLC3, FOXI1, PDE1C) and neuroendocrine cells (CHGA, CHGB, SCG3) in bat nasal organoids (Figure 2D; Supplemental Figure 3E). Intriguingly, we also observed a cell cluster in the bat distal lung organoids expressing both ciliated cell (FOXJ1, SNTN) and microfold cell markers (SPIB, SOX8, TNFAIP2) (Figure 2H; Supplemental Figure S2D). Microfold cells are antigen-sampling cells found in innate lymphoid tissue of the nasal (NALT) and intestinal tract (Peyer’s Patch), that use the RANKL/RANK signaling axis for maintenance and expansion^56, 57^ (Figure 2I). In the lungs of mice, microfold cells are present at a frequency of ∼0.08%, making them the rarest epithelial cell type found so far^58, 59^. In our bat distal lung organoids, ∼0.23% of the cells expressed microfold marker genes such as the bacterial uptake anchor protein GP2, the master regulators of microfold cell fate SOX8 and SPIB, as well as RANK (encoded by *TNFSRF11A*) and the RANK decoy receptor OPG (encoded by *TNFSRF11B*) (Figure 2J, K; Supplemental Figure S3E). Expression of the ligand to RANK (RANKL encoded by *TNFSF11*) was largely restricted to a subset of basal cells (Supplemental Figure S3F). Differential gene expression analysis further demonstrated specific transcriptional cell states in these rare cell types, including well-known markers and several novel candidate genes that warrant further exploration (Figure 2L). Our data shows that adult airway stem cell derived organoids from the Egyptian fruit bat recapitulate the complex cell type diversity of the stereotypical mammalian upper and lower airways, including the presence of rare specialized cell types important for mucosal immunity that are either absent or infrequently observed in single-cell RNA-sequencing data of human airway organoid cultures^60^. Our novel resource should allow researchers to explore bat airway epithelial cell states at unprecedented detail.

### Egyptian fruit bats small intestinal organoids resolved at single cell resolution

We further derived organoids from frozen small intestinal tissue of *Rousettus aegyptiacus* bats and were able to maintain them over at least six months in a niche-inspired human organoid medium^61^. Expanding gut organoids exhibited budded morphology with clearly visible interspersed granulated cells (Figure 3A). As previously shown for other bat species^33–35^, small intestinal organoids require activation of EGFR, FGFR and the WNT signaling pathways for growth and self-renewal (Supplemental Figure S4A). Removal of WNT3A and Noggin induced differentiation of KRT20+ and SLC2A2+ enterocytes in the bat organoids (Supplemental Figure S4B). In contrast, the intestinal stem cell marker LGR5 was significantly reduced in differentiated compared to expanding small intestinal organoids (Supplemental Figure S4B), demonstrating the need for WNT pathway activation to maintain Egyptian fruit bat intestinal stem cells *in vitro*.

**Figure 3.**
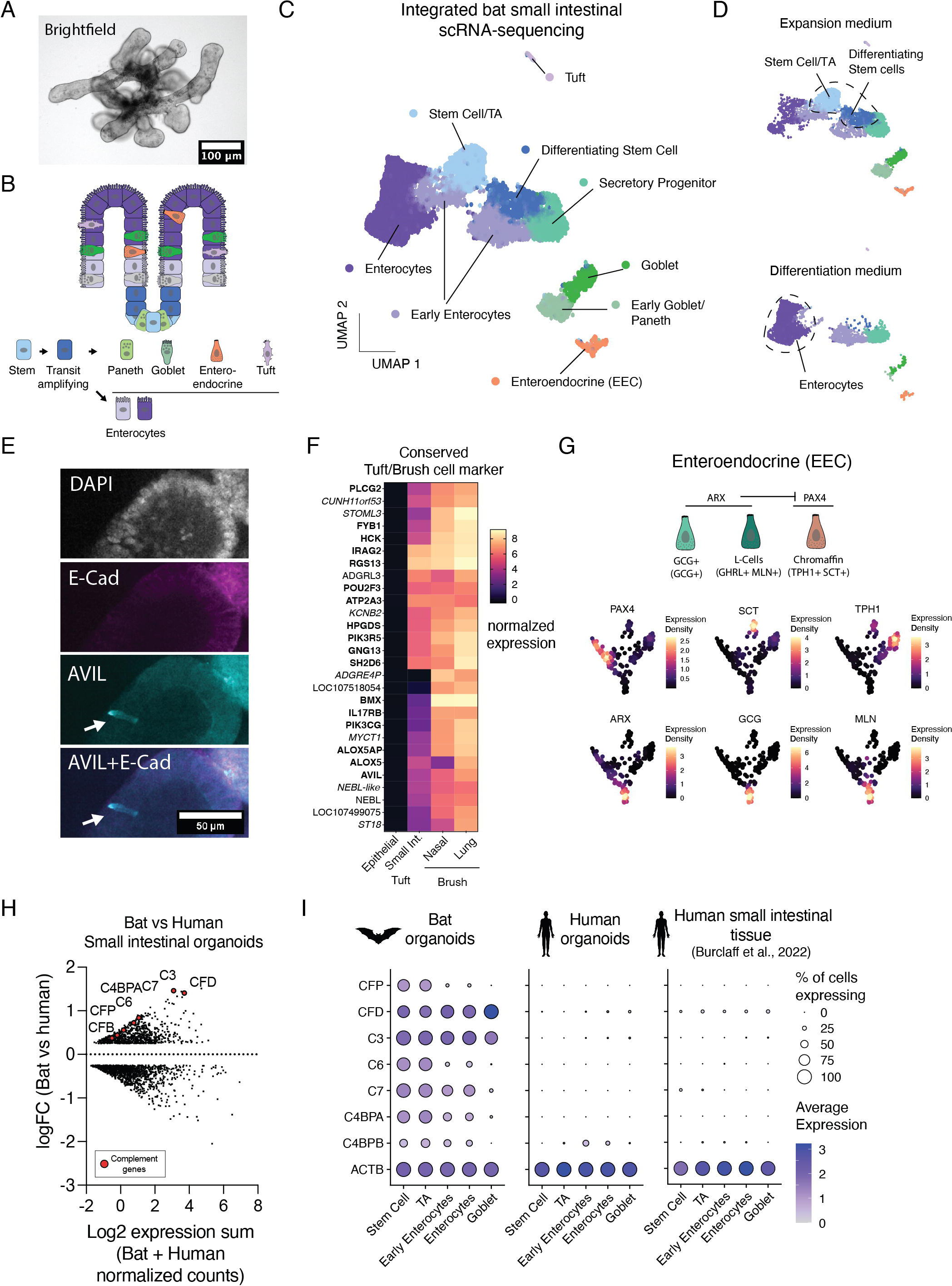
Single-cell RNA sequencing of bat small intestinal organoids. A) Brightfield image of bat small intestinal organoids. B) Schematic of cell type diversity found in the small intestinal epithelium. C) UMAP plot of integrated single-cell RNA sequencing dataset showing cell type clusters in bat small intestinal organoids. D) UMAP plot of integrated dataset split for the two different culture media conditions. Cell types showing major differences in abundance are shown. E) Immunofluorescence staining of tuft cells in small intestinal organoids. Anti-E-cadherin, Anti-Advilin (AVIL), DAPI (nuclei) and a merged representation are shown. The arrow indicates the solitary tuft cells expressing AVIL. F) A heatmap showing the average normalized expression values of conserved genes observed in intestinal tuft and airway brush cells. The three groups of tuft/brush cells are shown in comparison to an average of all other epithelial cell types. G) Schematic and expression of markers gene in enteroendocrine sub-lineages. Expression densities (Nebulosa package) for each marker gene are shown for enteroendocrine cells in UMAP space. The expression density gradient is depicted in the respective subfigure legend. H) Differential gene expression analysis comparing bat to human small intestinal organoids cultured in expansion medium. Each dot represents the log2 transformed SCT count normalized expression sum of a given gene from human and bat small intestinal organoids, and the log-fold (logFC) change from the analysis between the two species. Positive logFC values means upregulated in bat organoids compared to human organoids. Complement system genes are highlighted. I) DotPlot expression analysis showing the average expression of selected complement genes for bat or human small intestinal organoids, or external human small intestinal tissue^68^ determined from single-cell RNA sequencing data. The dot size shows the percentage of cells expressing a respective gene and the color shows the average expression level.

To profile the cell type composition, we performed single-cell RNA-sequencing of *Rousettus aegyptiacus* small intestinal organoids cultured in expansion or differentiation medium (Figure 3B; Supplemental Table 2). Single-cell RNA-sequencing analysis revealed the presence of intestinal stem cells (LGR5, SMOC2, CD44), transit amplifying cells (HELLS, PCNA), a cell population expressing early secretory and paneth cell markers (PLK2, SOX4, ATOH1, DEFA5, TFF3), goblet cells (MUC2, SPINK4), early enterocytes (KRT19-high, FABP5) and mature enterocytes (SLC2A2, ABCG2 (encoded by LOC107518949), KRT20) (Figure 3C; Supplemental Figure S4C). While differentiated bat organoids displayed a low number of intestinal stem cells compared to expanding organoids, they were strongly enriched in mature enterocytes (Figure 3D; Supplemental Figure S4D). Of note, the Coronavirus entry factors ACE2 (SARS-CoV/SARS-CoV-2)^62–64^ and DPP4 (MERS-CoV)^65^ were significantly upregulated upon organoid differentiation (Supplemental Figure S4E). Single-cell RNA-sequencing analyses also uncovered two distinct rare epithelial cell types, namely intestinal tuft (POU2F3, AVIL) and enteroendocrine cells (CHGA, CHGB) (Figure 3C, E; Supplemental Figure 4F). Interestingly, the bat intestinal tuft cells shared a set of markers with related airway brush cells (AVIL, POU2F3, PLCG2, RGS13, IRAG2, ALOX5AP, SH2D6, HCK, IL17RB, PIK3CG, BMX, SH2D6)^66^ (Figure 3F). Detailed assessment of enteroendocrine cells (CHGA+CHGB+) further revealed sub-lineages representing enterochromaffin cells (TPH1+SCT+), ARX-dependent GCG+ cells and MLN+ L-Cells^67^ (Figure 3G).

Comparative analysis of bat and human small intestinal organoids, cultured side-by-side, and previously published primary human small intestine tissue^68^ showed a strongly conserved transcriptional state for differentiated cells, except for paneth cells, which co-clustered with small intestinal tissue microfold cells (Supplemental Figure S5A-C). Bat organoids contained no microfold cells (GP2, SPIB), and only a few cells expressed known paneth cell markers (DEFA5, OTOP2, CA7), while human organoids were completely devoid of both (Supplemental Figure S5D). This is in line with previous studies showing that paneth and microfold cells are typically absent in human gut organoids unless cytokines such as IL-22 or RANKL are provided^69, 70^. Intriguingly, we noticed striking differences in the expression of genes from the classical and alternative complement systems comparing human and bat small intestinal organoids (Figure 3H; Supplemental Figure S6A-C). In contrast to human small intestinal organoids, bat organoids showed high expression of several complement genes (CFP, CFB, CFD, CFI, C2, C3, C4A (LOC107498435), C6, C7, C9, C4BPA, and CFH (LOC107520841)) (Figure 3H; Supplemental Figure S6A-C). Two inhibitors of complement activity (CD46, CD59) were strongly expressed in human but not bat organoids (Supplemental Figure S6A-C). Importantly, these expression differences were also apparent when comparing bat and human organoids to human small intestinal tissue using publicly available single-cell RNA-sequencing data^68^ (Figure 3I, Supplemental Figure S6D). These findings are in agreement with a recent pre-print study focusing on single-cell transcriptomics of *Rousettus aegyptiacus* small intestine and lung tissue^71^. Given that bat and human organoids were cultured under identical conditions, these differences are likely species-specific and warrant functional exploration. Collectively, we provide the first deep profiling data of bat small intestinal organoids and demonstrate that *Rousettus aegyptiacus* small intestinal organoids largely recapitulate the epithelial cell type diversity found in mammals, but also exhibit unexpected gene expression differences compared to humans.

### Antiviral responses to human-pathogenic Betacoronaviruses in bat organoids

Emerging RNA viruses utilize host factors for entry that often are exclusively expressed in differentiated cell types and absent in 2D cell culture. Thus, bat organoids hold the potential to better model natural infection scenarios without the need for ectopic expression of entry factors or live animals. To explore potential susceptibility of bat organoids to emerging viruses, we examined mRNA levels of known virus entry receptors in our single-cell RNA-sequencing datasets from basal cell derived nasal and distal lung air liquid interface cultures, alveolar organoids and small intestinal organoids. We found expression of several key entry factors with either broad (e.g Marburg/Ebola virus entry factor NPC1^72^) or more cell-type restricted expression patterns (e.g, ACE2 employed by SARS-CoV^62^/SARS-CoV-2^63^/NL63^73^ and NCAM1 used by RABV^74^) (Figure 4A). Thus, *Rousettus aegyptiacus* bat organoids may serve as infection models to cultivate and study bat-borne emerging viruses of human importance.

**Figure 4.**
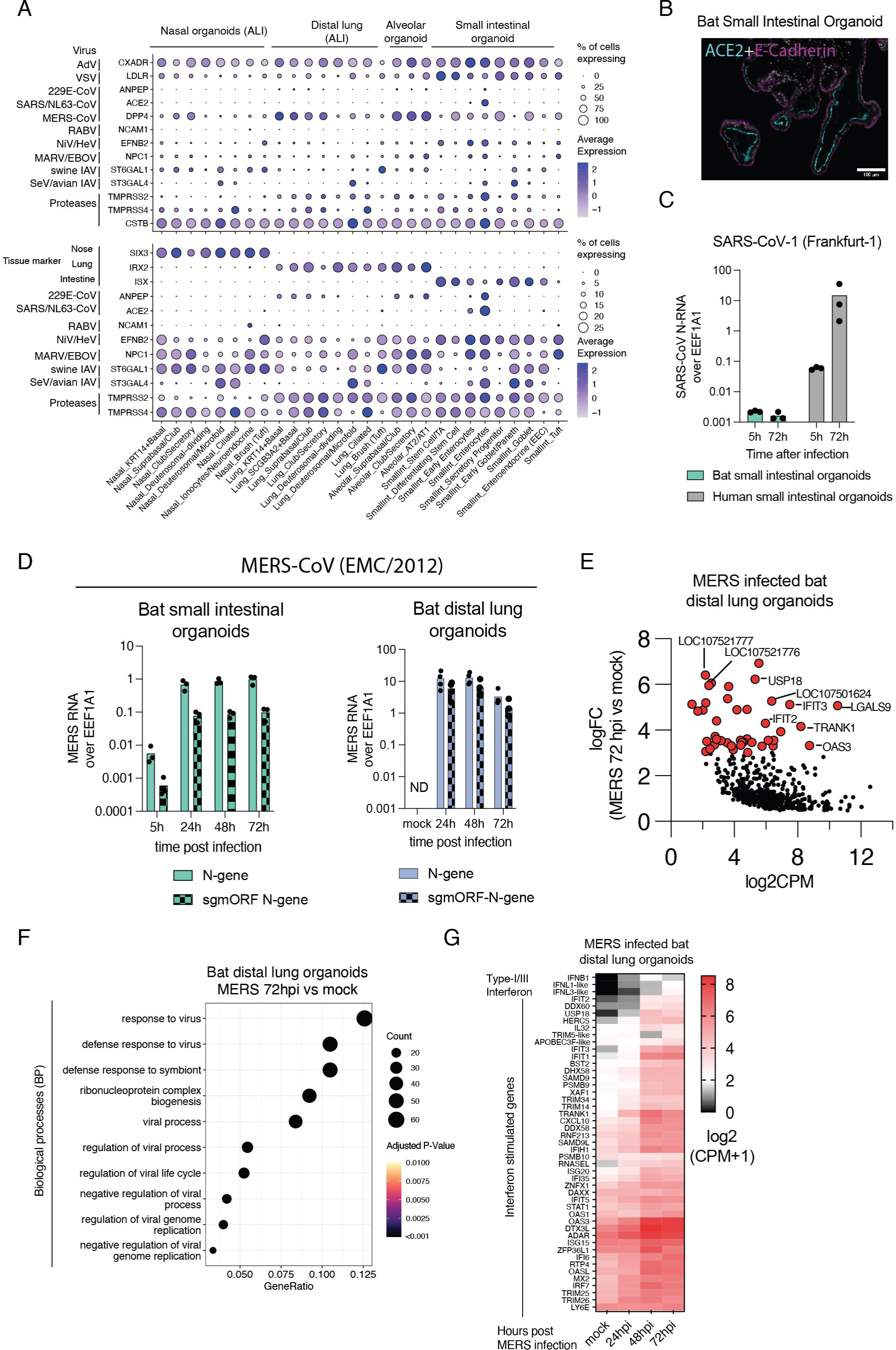
Infection of bat organoids with emerging Coronaviruses. A) DotPlot expression analysis showing the average expression of virus entry genes for each bat organoid cell type determined from single-cell RNA sequencing data. The dot size shows the percentage of cells in each cell type expressing a respective gene. The bottom panel shows a selected list of genes with the maximum percentage of cells expressing the feature (dot size) set to 25%. Genes expressed by more than 25% of cells have the same dot size. B) Immunofluorescence staining with anti-ACE2 and anti-E-Cadherin antibody in bat small intestinal organoids cultured in differentiation medium. C) RT-qPCR expression analysis of SARS-CoV-1 nucleocapsid gene in bat or human small intestinal organoids infected with SARS-CoV-1 at an MOI of ∼1 (50.000 PFUs) for 5 or 72 hours. Reference gene (EEF1A1) normalized expression values for cellular SARS-CoV-1 Nucleocapsid RNA are shown. Each dot represents an infected sample. D) RT-qPCR expression analysis of MERS Nucleocapsid RNA in bat small intestinal (left) or distal lung (right) organoids infected with MERS-CoV at an MOI of ∼1 (50.000 PFUs) for different times. Reference gene normalized expression values for cellular MERS nucleocapsid RNA or subgenomic RNA (leader-nucleocapsid gene) or are shown. Each dot represents an infected sample. E) Differential gene expression analysis of bat distal lung organoids infected with MERS for 72 hours or mock treated. Each dot represents the expression and log-fold change (logFC) of a differentially expressed gene between infected and uninfected organoid samples. Positive LogFC values mean upregulated in MERS infected organoids. Genes with a logFC > 3 are highlighted. F) Related to D). Biological process enrichment analysis performed with *clusterProfiler* of significantly upregulated genes at 72 hours post infection compared to uninfected cells. The top 10 enriched biological processes are shown. The values on the x-axis are the ratio of enriched to total genes for each biological process, while the dot size shows the number of enriched genes. The color shows the adjusted P-value. G) Heatmap showing the average log2 normalized RNA expression (EdgeR counts per million CPM+1) for interferons and interferon stimulated genes (ISG) in distal lung organoids following infection with MERS. Each value presents the average of three replicates after different hours post infection and mock infected.

In proof-of-concept experiments, we challenged *Rousettus aegyptiacus* organoids with emerging RNA viruses to test their susceptibility and investigate antiviral responses. We decided to focus on human viruses with confirmed or suspected bat origin, that can bind the cognate *Rousettus aegyptiacus* receptors, have documented spillover events and cause substantial morbidity in humans. Given the specific and high expression of ACE2 in intestinal enterocytes (Figure 4A, B), we first infected bat and human small intestinal organoids with the human Coronavirus SARS-CoV-1 (human case fatality rate up to 10%^75^), which was shown to efficiently bind human and *Rousettus aegyptiacus* ACE2 in pseudotype virus entry assays^76^. Unlike human small intestinal organoids which were readily infected in parallel experiments, we failed to observe any apparent SARS-CoV-1 replication in bat organoids (Figure 4C). Similar findings have been reported using infections with the SARS-CoV-1 like Coronavirus WIV1, which was unable to replicate in Egyptian fruit bats despite efficient binding of WIV1 spike glycoprotein to bat ACE2^77^.

We next challenged bat small intestinal and distal lung organoids displaying high DPP4 expression with the human betacoronavirus MERS-CoV (EMC/2012), which causes severe lung disease and death in up to 35% of infected humans^78^. We observed infection and replication of MERS-CoV in alveolar and small intestinal organoids by examining genomic and subgenomic RNA expression with RT-qPCR (Figure 4D; Supplemental Figure S7A). Viral genomic and subgenomic RNA was at least 10-fold higher in alveolar compared to intestinal organoids, consistent with the highest expression of DPP4 observed in alveolar AT2 cells (Figure 4A, D; Supplemental Figure S7B), which is also the major target cell type for MERS-CoV in humans^79^. Differential gene expression analyses of MERS-CoV infected distal lung bat organoids revealed a strong induction of antiviral defense genes, including type-I/III interferons (IFNB1, IFNL1-like (LOC107521777), IFNL3-like (LOC107521776)) and interferon stimulated genes (ISGs) involved in virus sensing (TLR3, DDX58, IFIH1), virus restriction (IFIT1/2/3, IFITM1, OAS1/3, MX1/2, ADAR/APOBEC3-like, BST2), or host cell antiviral signaling (SOCS1, USP18, IRF7, STAT1) (Figure 4E; Supplemental Table 3). GO-term analysis confirmed a strong enrichment of biological processes related to antiviral defense and interferon signaling upon MERS-CoV infection (Figure 4F). Expression of Interferon and ISGs were dynamically regulated, increasing in RNA levels throughout the infection time course (Figure 4G). Of note, our results in bat organoids are distinct to MERS-CoV infections in human cell lines and differentiated primary airway models reported previously^80^. These data show that lung organoids of Egyptian fruit bats support MERS-CoV replication and respond to infection with a type-I/III interferon driven antiviral immune response.

### Marburg virus infections in bat and human organoids

We next studied Marburg virus infections in our *Rousettus aegyptiacus* bat organoids. Egyptian fruit bats are the confirmed natural reservoir for MARV and remain asymptomatic following natural or experimental infection^7, 14^. In humans, Marburg virus infections can induce a fatal hemorrhagic fever disease with case fatality rates up to 90%^81^. In agreement with the broad cellular expression of the MARV entry receptor NPC1, we found productive replication of MARV (Musoke isolate) by RT-PCR in all tested bat organoids, including bat nasal air-liquid interface (ALI) cultures, alveolar and small intestinal organoids (Figure 4A, 5A). MARV infected human bronchial epithelial cells grown at the air liquid interface also displayed an increase in MARV L-gene RNA expression over time, similar to the corresponding bat airway organoid cultures (Figure 5A). These data demonstrate that Egyptian fruit bat organoids and differentiated human airway cultures support entry and replication of MARV.

**Figure 5.**
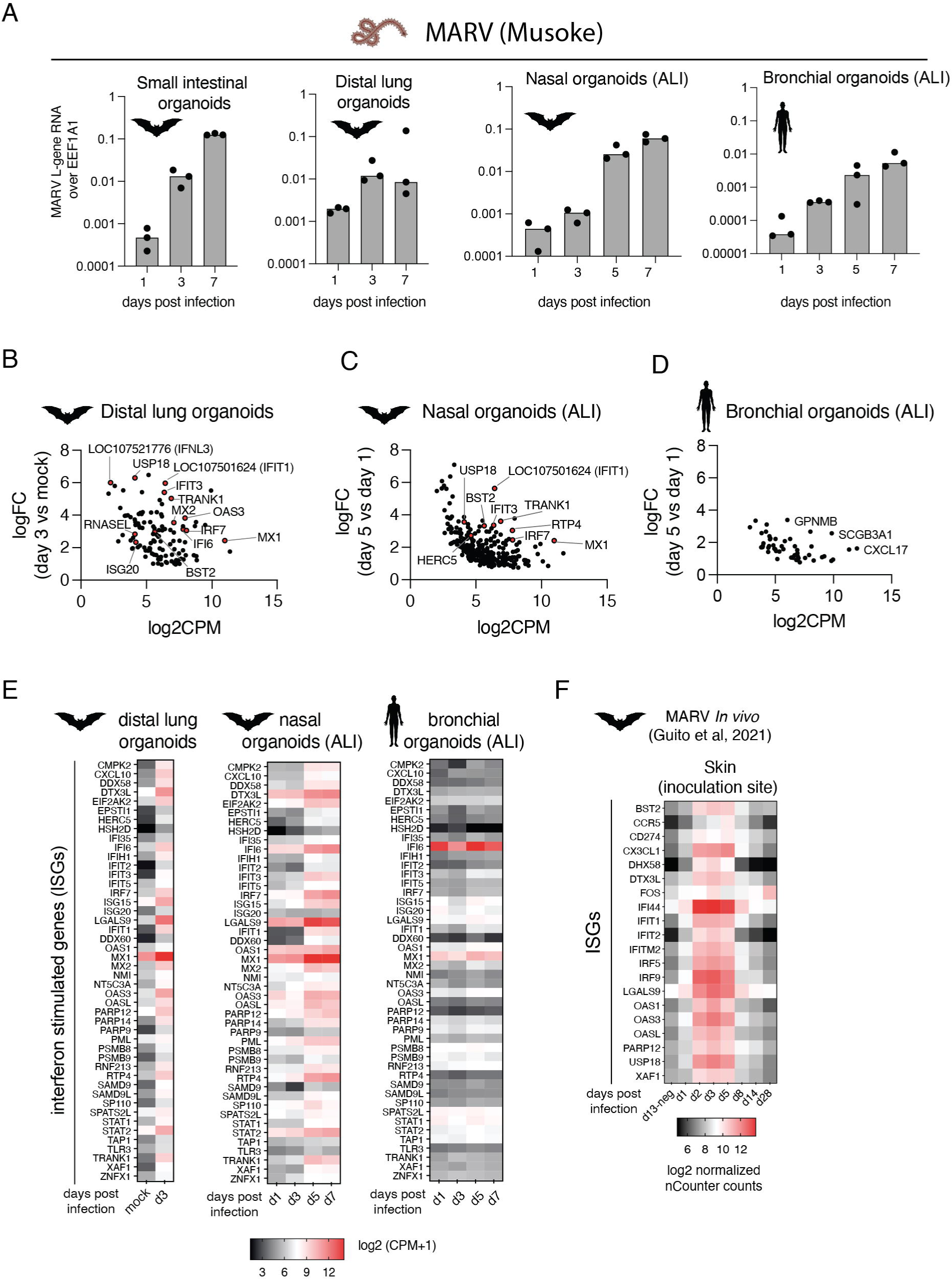
Infection of bat organoids with Marburg virus. A) RT-qPCR expression analysis of MARV L-gene RNA in different organoid models infected with MARV at an MOI of 0.5-1 (50.000 PFUs, or 100.000 PFU for ALI) for different times. Reference gene normalized expression values for MARV L-gene RNA are shown. Each dot represents an infected sample. B) Differential gene expression analysis of bat distal lung organoids infected with MARV for 72 hours. Each dot represents the expression and log-fold change of a differentially expressed gene between infected and uninfected organoid samples. Selected genes are highlighted. C) Differential gene expression analysis of bat nasal ALI cultures infected with MARV for 5 days compared day 1 infected cultures. Each dot represents the expression and log-fold change of a differentially expressed gene between day 5 and day 1 infected samples. Selected genes are highlighted. D) Differential gene expression analysis of human bronchial ALI cultures infected with MARV for 5 days compared day 1 infected cultures. Each dot represents the expression and log-fold change of a differentially expressed gene between day 5 and day 1 infected samples. Selected genes are highlighted. E) Heatmap showing log2 normalized gene expression (log2 counts per million + 1) for interferon stimulated genes (ISGs) in uninfected and MARV infected bat distal lung organoids (left), MARV infected bat nasal ALI cultures (middle), MARV infected human bronchial ALI cultures (right). F) Data from *In vivo* MARV infection of Egyptian fruit bats (from Guito et al, 2021): Log2-transformed nCounter normalized expression of selected, dynamically regulated ISGs at the virus inoculation site (skin tissue) throughout the infection time-course.

Aerosolized MARV causes lethal infection in animals^82–84^, supporting an entry route via the respiratory system. We therefore decided to further investigate transcriptional changes upon MARV infection in bat and human airway organoid cultures. Gene expression profiling showed a significant induction of antiviral response genes in bat distal lung and bat nasal organoids differentiated at the air-liquid interface following MARV infections (Figure 5B, C, E; Supplemental Table 3). Gene-ontology enrichment analysis of infected bat organoids further revealed induction of multiple pathways related to virus defense and interferon signaling (Supplemental Figure S7C, D). These data demonstrate a strong antiviral innate immune response in Egyptian fruit bat upper and lower airway epithelial cells following MARV infection *in vitro*. In contrast, MARV infected human bronchial air-liquid-interface cultures showed little to no induction of interferon regulated genes (Figure 5D, E), despite these differentiated human epithelial cells potently inducing an antiviral response following Sendai virus (also known as murine respirovirus) infection (Supplemental Figure S7E, F; Supplemental Table 3). To compare our findings to previous results obtained from *in vivo* MARV infection data of Egyptian fruit bats, we examined published gene expression data^7^. Like our organoid infection data, Egyptian fruit bats display a strong and temporally regulated induction of interferon stimulated genes at the site of infection, and to an extent in other organs even when viral levels were low (Figure 5F; Supplemental Figure S8). These data show that bat respiratory organoids exhibit strong innate antiviral immune responses to Marburg virus that are blunted in differentiated human airway epithelial cell cultures.

### Expression of interferon genes in Egyptian fruit bat organoids

Our infection data clearly showed that emerging RNA virus infections unleash an antiviral immune response in bat organoids, which is characterized by the induction of genes of the type-I and type-III interferon system. Although type-III interferons (IFN-λ) orchestrate antiviral defenses at infected epithelial surfaces^85^, they have received little attention in bats^86^. We therefore sought to decode responses to exogenous interferon-α (type-I interferon) and interferon-λ (type-III interferon) in bat and human epithelial organoids. Basal expression of interferon genes has been described for *Pteropodidae* bats^23^, presenting a potential confounder in interferon stimulation experiments. We therefore first examined if any interferon genes are constitutively expressed in our bat organoid single-cell RNA-sequencing dataset, finding no apparent RNA expression of canonical type-I/II/III interferons in airway and small intestinal bat organoids (Figure 6A).

**Figure 6.**
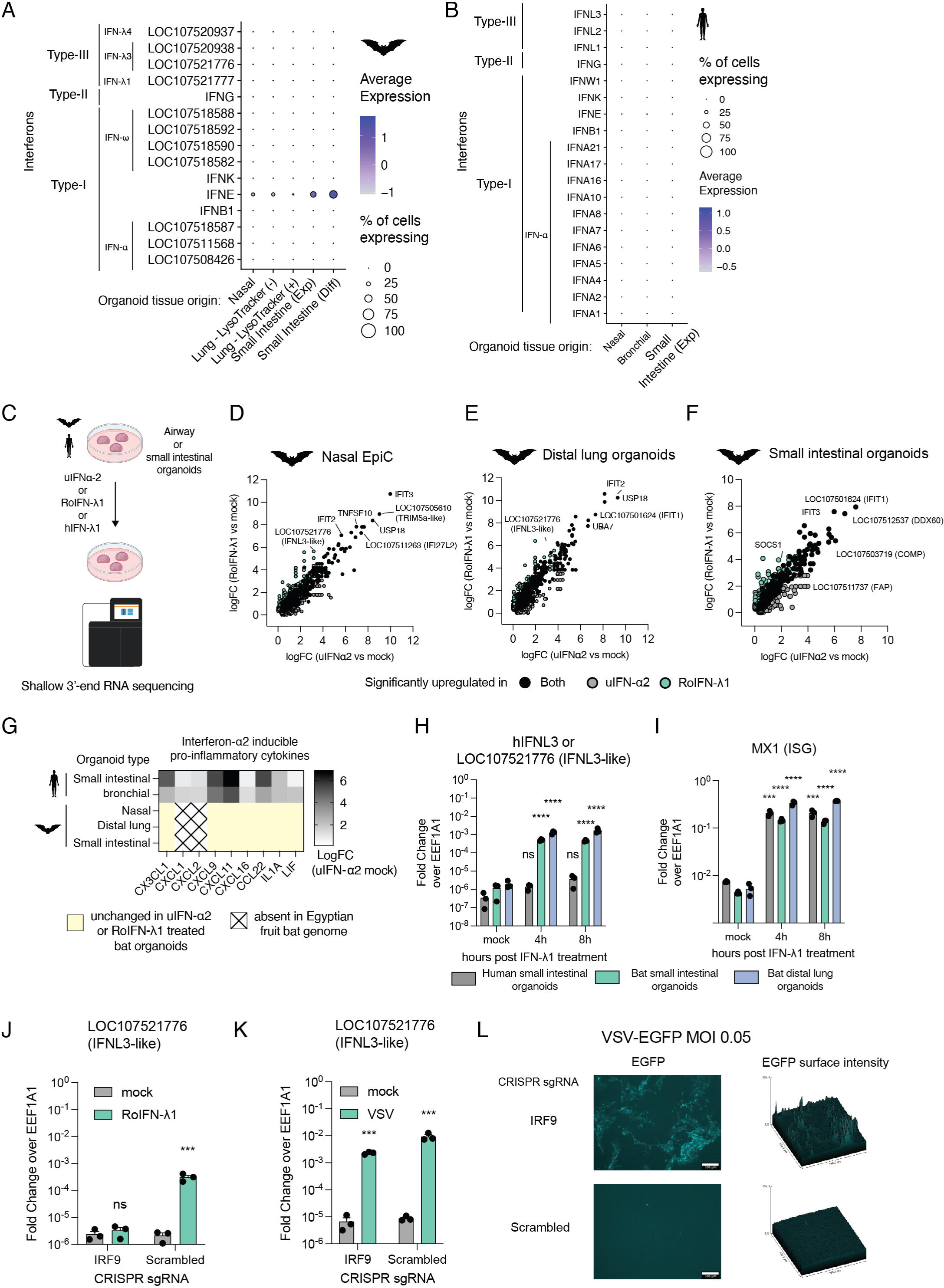
Decoding the bat epithelial interferon response in organoids. A) DotPlot expression analysis showing the average expression of interferon genes for each bat organoid systems determined from single-cell RNA sequencing data. The dot size shows the percentage of cells in expressing a respective gene and the color shows the average expression level. B) DotPlot expression analysis showing the average expression of interferon genes for each human organoid systems determined from single-cell RNA sequencing data. The dot size shows the percentage of cells in expressing a respective gene and the color shows the average expression level. C) Schematic of experimental overview involving interferon stimulation and readout by RNA-sequencing. D) Scatterplot showing the log-fold changes of genes determined by comparing the expression of interferon to mock treated nasal organoid derived primary epithelial cells (Nasal EpiC). Each dot represents a pairwise-comparison in either IFNα2 to mock or RoIFN-λ1 to mock treated cells. Genes significantly upregulated after both uIFN-α2 and IFN-λ1 interferon stimulation or only by either one are colorcoded. Individual genes of interest are highlighted. E) Same as in E) but for bat distal lung organoids. F) Same as in E) but for bat small intestinal organoids. G) Heatmap showing log-fold changes (logFC) for a selected list of pro-inflammatory cytokines in interferon-α2 versus mock treated human or bat organoids. If there was no observed expression change (e.g in bat organoids), a pseudovalue of 0 was given and highlighted in yellow. H) RT-qPCR expression analysis of human IFNL3 or bat IFNL3-like in different organoids at different time-points following RoIFN-λ1 or hIFN-λ1 treatment. EEF1A1 reference gene normalized expression values for IFNL3 are shown. Each dot represents a biological replicate. Unpaired student t-tests were performed between mock and treatments (ns: not significant, * P<0.05, ** P<0.01, *** P<0.001, **** P<0.0001). I) Same as in H) but for MX1. J) RT-qPCR expression analysis of bat IFNL3-like in bat small intestinal organoids expressing Cas9 and a scrambled guide RNAs treated RoIFN-λ1 for eight hours. EEF1A1 reference gene normalized expression values for bat IFNL3-like are shown. Each dot represents a biological replicate. Unpaired student t-tests were performed between mock and treatments (ns: not significant, * P<0.05, ** P<0.01, *** P<0.001, **** P<0.0001). K) RT-qPCR expression analysis of bat IFNL3-like in differentiated bat small intestinal organoids expressing Cas9 and a scrambled control guide RNA infected with VSV-G at an MOI of 0.05 for 72 hours. EEF1A1 reference gene normalized expression values for IFNL3-like are shown. Each dot represents a biological replicate. Unpaired student t-tests were performed between mock and infections (ns: not significant, * P<0.05, ** P<0.01, *** P<0.001, **** P<0.0001). L) EGFP fluorescence image of organoids expressing Cas9 and different guide RNAs infected with VSV-GFP for 72 hours. The EGFP surface intensity plot (ImageJ) of the image is shown on the right.

Surprisingly, and unlike in human organoids, we found high constitutive expression of interferon-ε (IFNE) in defined bat organoid cell types, especially in small intestinal enterocytes and nasal KRT5/KRT14+ basal cells (Figure 6A, B; Supplemental Figure S9A). Interferon-ε (IFNE) is a non-canonical type-I interferon constitutively expressed in the female reproductive tract of mice and humans, protecting these tissues from viral and bacterial infections^87^. Using publicly available *in vivo* tissue gene expression data of Egyptian fruit bats (GSE152728)^88^, we could also identify interferon-ε expression in several tissues, such as lung, salivary gland, kidney, and, showing the highest expression, in the small intestine; with little to no IFNE transcript counts in bat liver and peripheral blood mononuclear cells (PBMC) (Supplemental Figure S9B). Immunofluorescence staining in *Rousettus aegyptiacus* revealed strong interferon-ε immunoreactivity in bat small intestinal tissue and differentiated bat small intestinal organoids (Supplemental Figure S9C, D). Anti-Interferon-ε reactivity was weak or absent in liver tissue and isotype-control stains (Supplemental Figure S9C). Thus, except for interferon-ε, type-I or type-III interferon genes are not constitutively expressed in Egyptian fruit bats *in vitro* and *in vivo*^22^.

### Decoding Egyptian fruit bat epithelial type-I/type-III interferon responses

To further delineate interferon responses in *Rousettus aegypticaus*, we decided to stimulate bat distal lung and small intestinal organoids, as well as nasal organoid derived basal epithelial cells with type-I and type III interferons and monitor gene expression with 3’-end bulk RNA-sequencing (Figure 6C). Commercially available universal Interferon-α2 was used to monitor type I interferon activation, serving as positive control and point of comparison between type-I and type-III interferon responses. To reveal possible species-specific differences between bats and humans, we also decided to stimulate human small intestinal and bronchial organoids with recombinant universal Interferon-α2 and human IFN-λ1. Bat type-III interferons are not commercially available, which required us to produce recombinant *Rousettus aegypticaus* IFN-λ in human HEK293T cells. For initial experiments, we focused on interferon-λ1-like (encoded by LOC107521777), an orthologue of human IFNL1. Bat interferon-λ1-like (hereafter termed RoIFN-λ1) was induced in bat lung and small intestinal organoids infected with emerging viruses or Sendai virus, a commonly used RNA virus known to stimulate the RNA virus pattern recognition receptors RIG-I^89^ and MDA5^90^ (IFIH1) (Supplemental Figure S10A, B). RoIFN-λ1 as well as universal Interferon-α2 exhibited a potent dose-dependent activation of the interferon stimulated gene IFIT1 (encoded by LOC107501624) (Supplemental Figure S10C, D). Combined addition of interferon and the JAK1/2 inhibitor Ruxolitinib prevented the induction of the interferon-stimulated genes ISG15, IFIT3, measured by RT-qPCR (Supplemental Figure S10E). These data demonstrate that both universal interferon-α2 and, importantly, bat RoIFN-λ1 activation signals are mediated via the JAK/STAT axis in primary bat epithelial cells.

Genome-wide mRNA expression profiling in bat organoids showed that interferon-α2 and RoIFN-λ1 induce an overlapping set of genes (Figure 6D-F; Supplemental Table 4). Of note, the number of differentially upregulated genes in bat organoids was consistently greater in response to RoIFN-λ1 stimulation compared to universal interferon-α2 (Supplemental Figure S10F;). Antiviral defense and interferon response gene ontology-terms were the top enriched biological processes in response to RoIFN-λ1 and universal interferon-α2 treatment (Supplemental Figure S10G, H). Approximately one hundred genes were significantly upregulated across all conditions, including several known interferon-regulated genes (e.g. ADAR, BST2, IFIT1/2/3, DDX60/DDX60L, MX1/2, ISG15, IRF7, USP18, OAS1/3/OASL, RTP4). Genes significantly upregulated by RoIFN-λ1 but not universal interferon-α2 in bat organoids included virus restriction factors (NCOA7, APOBEC-3C-like, HELZ2) and immune response regulators (ETV7, PSME1). Human small intestinal and bronchial organoid air-liquid interface cultures showed a similarly positive correlation in gene expression changes upon human IFN-λ1 and universal interferon-α2 treatment (Supplemental Figure S11A-D; Supplemental Table 4). Of note, universal IFN-α2 upregulated genes were greater in number compared to hIFN-λ1 stimulated genes in human organoid cultures (Supplemental Figure S11E). We also noted a stronger induction of pro-inflammatory chemokines (e.g CXCL1, CXCL2, CXCL9, CXCL11, CX3CL1, CCL22) in universal interferon-α2 compared to IFN-λ1 stimulated human respiratory and intestinal organoids (Supplemental Figure S11A, B). By contrast, bat organoids displayed no or minimal expression of these pro-inflammatory genes after universal interferon-α2 or bat RoIFN-λ1 treatment (Figure 6G). This data show that unlike commonly used bat *in vitro* cell lines, bat epithelial organoids not only produce type-III interferons upon virus infection, but these interferons can induce a potent antiviral gene expression state.

### IFN-λ3 is an interferon-regulated gene in Egyptian fruit bats

Our results surprisingly revealed a strong induction of the putative interferon-λ3-like (LOC107521776) gene in RoIFN-λ1 treated bat organoid models. Using RT-qPCR, we confirmed that interferon-λ3-like was significantly upregulated early after interferon-λ1 stimulation in bat small intestinal and distal lung, but not human small intestinal organoids, despite an equally strong induction of the interferon-stimulated gene MX1 (Figure 6H, I). To confirm the functional role of the putative interferon-λ3-like protein, we produced bat recombinant RoIFN-λ3-like protein (encoded by LOC107521776 and LOC107520938) (Supplemental Figure S12A). Like interferon-λ1-like, bat interferon-λ3-like stimulation of bat small intestinal organoids from *Rousettus aegyptiacus* led to a significant upregulation of multiple known interferon stimulated genes such as MX1/2, OAS3, or IFIT1/2/3 (Supplemental Figure S12B-E; Supplemental Table 4). Consistent with induced antiviral transcriptional states induced by interferon-λ1 or interferon-λ3-like proteins, pre-treatment of bat nasal, tracheal, and small intestinal organoids with these cytokines conferred protection against VSV-EGFP virus (Supplemental Figure 13A, B). We further find that antiviral protection could be maintained in nasal organoid derived monolayer cultures in the absence of interferon stimulation for at least 12 hours (Supplemental Figure 13C-F).

To determine genetic dependencies, we established an efficient CRISPR-Cas9 genome editing system in bat organoids that utilizes stable integration of Cas9 and sgRNAs through Lentivirus delivery (Supplemental Figure S14A). We decided to focus on IRF9 because it is an integral transcription factor mediating signaling by both type-I and type-III interferons^91^, thus allowing us to determine if IFN-λ3 is indeed a canonical interferon stimulated gene. Interferon-λ1 mediated induction of interferon-λ3-like was indeed genetically dependent on IRF9, as CRISPR-Cas9 perturbation of IRF9 in bat small intestinal organoids almost completely abrogated induction of the RoIFN-λ3-like gene (LOC107521776) as well as induction of the interferon-stimulated gene MX1 (Figure 6J, Supplemental Figure S14B). In contrast, genetic ablation of IRF9 did not prevent the induction of IFN-λ3-like, but abolished MX1 expression following infection with vesicular stomatitis virus encoding EGFP (VSV-EGFP), highlighting virus dependent and virus independent pathways of interferon-λ3-like induction (Figure 6K; Supplemental Figure S14C). Endogenous interferon signaling and protection from VSV-EGFP replication however was also visibly impaired in IRF9 knockout organoids (Figure 6L). Taken together, we provide the first comprehensive functional description of the Egyptian fruit bat type-III interferon system in respiratory and intestinal organoids *in vitro* and report a novel aspect of interferon-λ gene regulation, identifying a paradigmatic amplifying circuit of the type-III interferon responses in bats.

## Discussion

Bats host a large diversity of viruses without showing signs of clinical disease, even after experimental infection^6–8, 17, 38^. However, differences in viral disease etiology between bats and other mammals are complex, encompassing both primary responses at the site of infection and peripheral immune reactions. Therefore, tissue-like *in vitro* model systems are critically needed to dissect the molecular mechanisms of often asymptomatic virus infections in bats. To delineate innate immune responses in infected epithelial tissues, we therefore established self-replicating organoids from cryogenically preserved tissue of the upper and lower respiratory tract, and small intestinal epithelium of *Rousettus aegyptiacus* bats. These self-replicating organoids can be passaged, cryopreserved and are amenable to CRISPR-Cas9 genetic perturbations, providing a sustainable organoid platform to functionally probe the unique biology of bats and to democratize access to bat research.

We chose the Egyptian fruit bat for several reasons. First, *Rousettus aegyptiacus* bats naturally tolerate pathogenic viruses that can infect humans such as Marburg virus (MARV)^7^, Sosuga virus (SOSV)^38^, Kasokero virus (KASV)^39^, or Rousettus bat H9N2 Influenza-A virus^92^. Second, the Egyptian fruit bat genome is the one of the most contiguous bat genomes available^22^, facilitating gene annotations, in-depth gene expression analysis and genetic perturbation experiments. Moreover, combining the molecular and cellular data reported in our current study with previous *in vivo* and *in vitro* studies in Egyptian fruit bats with the access to breeding colonies^7, 22, 88, 93, 94^ should markedly advance experimental bat research. Thus, Egyptian fruit bats are and will continue to be a cornerstone bat species for future studies.

Our single-cell RNA profiling data demonstrates that like for human organoids^44, 60, 61, 95^, Egyptian fruit bat adult stem cell-derived organoids largely recapitulate *in vivo* epithelial cell type diversity. One exception were airway goblet cells, which were of low abundance but could be readily induced by IL-13 stimulation. These findings are consistent with a recent report on tracheal organoids from *Eonycteris spelaea*^36^ and *Carollia perspicillata*^37^, suggesting that external factors present *in vivo* control airway goblet cell formation and maintenance. Single-cell RNA profiling further identified rare cell types that include lung microfold cells (M-cells), which specialize in luminal antigen uptake and presentation to intraepithelial immune cells. Given the low percentage of microfold cells in the mouse lung (<0.1%), it remains poorly understood how these cells emerge. A recent study^58^ combined RANKL stimulation with lineage tracing in murine lung cultures, concluding that mouse lung M-cells likely originate from Scgb1a1 expressing club cells, the direct precursor of ciliated or goblet cells. In our work, bat lung M-cells emerged without exogenous RANKL stimulation, and closely clustered with ciliated cells and intermediate cells expressing key cell fate determinants of both cell types, i.e. FOXJ1^96^ in ciliated cells and SPIB^97^ in M-cells. This novel relationship to ciliated cells warrants further exploration, but we speculate that bat lung microfold cells may arise from a direct ciliated cell progenitor expressing FOXJ1. Thus, bat lung organoids provide an *in vitro* model to investigate “naturally” occurring microfold cell emergence and function.

Limited or absent expression of virus entry receptors in cell lines often necessitates the ectopic overexpression of these molecules to render naturally unsusceptible cells permissive to virus infections^20^. Single-cell transcriptional profiling in our respiratory and intestinal organoids showed expression of multiple viral entry receptors in 3D engineered bat tissues, making them amendable to model natural infection scenarios and study *bona fide* cellular responses to virus infections. For instance, our organoids express ACE2 and DPP4, the key entry receptors for various Betacoronaviruses. Interestingly, infection experiments revealed that SARS-CoV-1 was unable to infect Egyptian fruit bat organoids despite high levels of endogenous ACE2. Although previous studies have shown that SARS-CoV-1 replicates in immortalized Egyptian fruit bat RoNi/7 cells expressing human ACE2 ^98^, our data does not support a natural infection scenario via ACE2 expressing epithelial tissue in Egyptian fruit bats. These findings are in line with the inability of SARS-CoV-1-related Coronavirus WIV1 to propagate in Egyptian fruit bats^77^. By contrast, MERS replicated in distal lung bat organoids and induced a strong type-I/III interferon-mediated antiviral response; which occurs only to a limited extent in differentiated primary human airway epithelial cultures as previously reported^60, 80^. Although MERS has not been found in *Rousettus aegyptiacus*, the related group 2d Betacoronavirus HKU9 RNA has been detected in *Rousettus* bats in independent sampling studies^99, 100^. Thus, Egyptian fruit bat may serve as a suitable surrogate to study MERS coronavirus pathogenesis in bats compared to other naturally susceptible mammals including humans.

Contrary to previous observations from *in vitro* infection studies on immortalized Egyptian fruit bat cell lines^93, 94^, we observed a significant upregulation of interferon-stimulated genes in MARV infected bat airway organoids without apparant activation of pro-inflammatory pathways. In general, we did not observe overt inflammatory responses in virus infected or interferon treated bat organoids, although epithelial organoids may imperfectly model these immune reactions. Nevertheless, these data paralleled infection responses observed in MARV infected Egyptian fruit bat animals, especially those obtained from samples from the primary site of infection^7^. Interestingly, this effect was not seen in our MARV infected human airway cultures, consistent with previous reports on MARV infections in primary human^101^ and bat cells^102^. Asymptomatic MARV infection in Egyptian fruit bats could therefore be the result of strong induction of the innate antiviral immunity at infected barrier tissue restriciting virus dissemination, and concurrent limitation of inflammatory pathways. In contrast, the inability of humans and non-human primates to mount a potent early antiviral response may lead to the observed uncontrolled virus replication, systemic inflammation and lethal infections in these species^82, 103, 104^. A mechanistic explanation for the species-specific differences may be rooted in the ability of bats to overcome effective innate immune antagonisms encoded by filoviruses^103^ and other emerging pathogens^105^. Investigating host-pathogen interactions in bats and humans beyond entry requirements will therefore be of critical importance to understand the ecology, spillover potential and human pathogenesis of bat-borne viruses. Organoids can provide a viable platform to study molecular host-pathogen interactions in different species under defined conditions.

Although being investigated for over twenty years, our study on bat organoids has uncovered novel features of the bat interferon system. For example, we discovered constitutive expression of interferon-ε in bat but not human organoids. It is tempting to speculate that interferon-ε may contribute to a primed antiviral state in the respiratory and intestinal mucosal tissue of Egyptian fruit bats, providing greater resistance to viruses compared to other mammals including humans. Alternatively, in Egyptian fruit bats interferon-ε might play a role in tissue homeostasis independent of classical interferon signaling^106^. Our work also entails a first comprehensive functional comparison of type-I and type-III epithelial interferon responses in bats, the latter of which are rapidly and strongly induced after virus infections and can confer antiviral immunity in bat organoids. Species comparisons also revealed the induction of pro-inflammatory chemokines in interferon-α2 treated human but not bat organoids, mechanistically linking dysregulated interferon responses to immune pathologies observed in emerging viral diseases in humans^103, 107^. Importantly, we also discovered Egyptian fruit IFN-λ3-like as an interferon stimulated gene, which could drive amplification of interferon responses to locally restrict virus replication at the primary site of infection. Together, our data indicate that type-III interferon responses play an essential role in bat mucosal antiviral immunity which can now be further mechanistically explored using our organoid resources and possibly translated to humans.

In summary, *Rousettus aegyptiacus* organoids provide an invaluable tool to study antiviral immunity in asymptomatic reservoir species. Our comprehensive single-cell RNA-sequencing data of organoids from the entire respiratory epithelial tree and small intestine of Egytpian fruit bats, combined with detailed molecular examination of the epithelial innate immune response to interferons and multiple emerging viruses, establish a molecular framework to mechanistically dissect bat epithelial biology under controlled laboratory conditions. Our cellular and molecular resources pave the way for functional genetic and mechanistic research in complex cellular tissue models of bats. Importantly, our result uncover fundamental circuits and differences in innate immune responses to viruses that could explain why bats can serve as reservoir species for viruses of pandemic potential that are highly lethal in humans.

## Acknowledgments and Funding

We thank all members of the Penninger laboratory, Johannes Zuber (IMP, Vienna) and Sylvia Knapp (Medical University of Vienna) for their support and critical feedback on the presented work. We would also like to thank in-house and Vienna Biocenter core facilities (VBCF) for providing excellent scientific infrastructure and services, especially the BioOptics facility, Molecular Biology Service, the VBCF Histology facility, the Next Generation Sequencing facility (Thomas Grentzinger for preparing 10x scRNA-sequencing libraries) and Bioinformatic support (Maria Novatchkova for providing help regarding the *souporcell* pipeline).

The laboratory of J.M.P. received funding from the Austrian Academy of Sciences, the Medical University of Vienna, the Vienna Science and Technology Fund (WWTF; 10.47379/EICOV20002), the Swedish Research Council (<u>2018-05766</u>), the Fundacio La Marato de TV3 (<u>202125-31</u>), the T. von Zastrow foundation, the Canada 150 Research Chairs Program F18-01336, the Canadian Institutes of Health Research COVID-19 grants F20-02343 and F20-02015 and the Innovative Medicines Initiative 2 Joint Undertaking under grant agreement No <u>101005026</u>. This Joint Undertaking receives support from the European Union’s Horizon 2020 research and innovation programme and EFPIA. The authors would also like to thank the Leducq Foundation for supporting their research with the Transatlantic Network of Excellence grant “ReVAMP — Recalibrating Mechanotransduction in Vascular Malformations” (21CVD03) and the Diamond-Blackfan-Anemia Fundraising (dbaexperiment.org). We also gratefully acknowledge funding by the German Federal Ministry of Education and Research (BMBF) under the project “Microbial Stargazing – Erforschung von Resilienzmechanismen von Mikroben und Menschen” (Ref. 01KX2324).

## Legends to Supplemental Figures

**Supplemental Figure S1.**
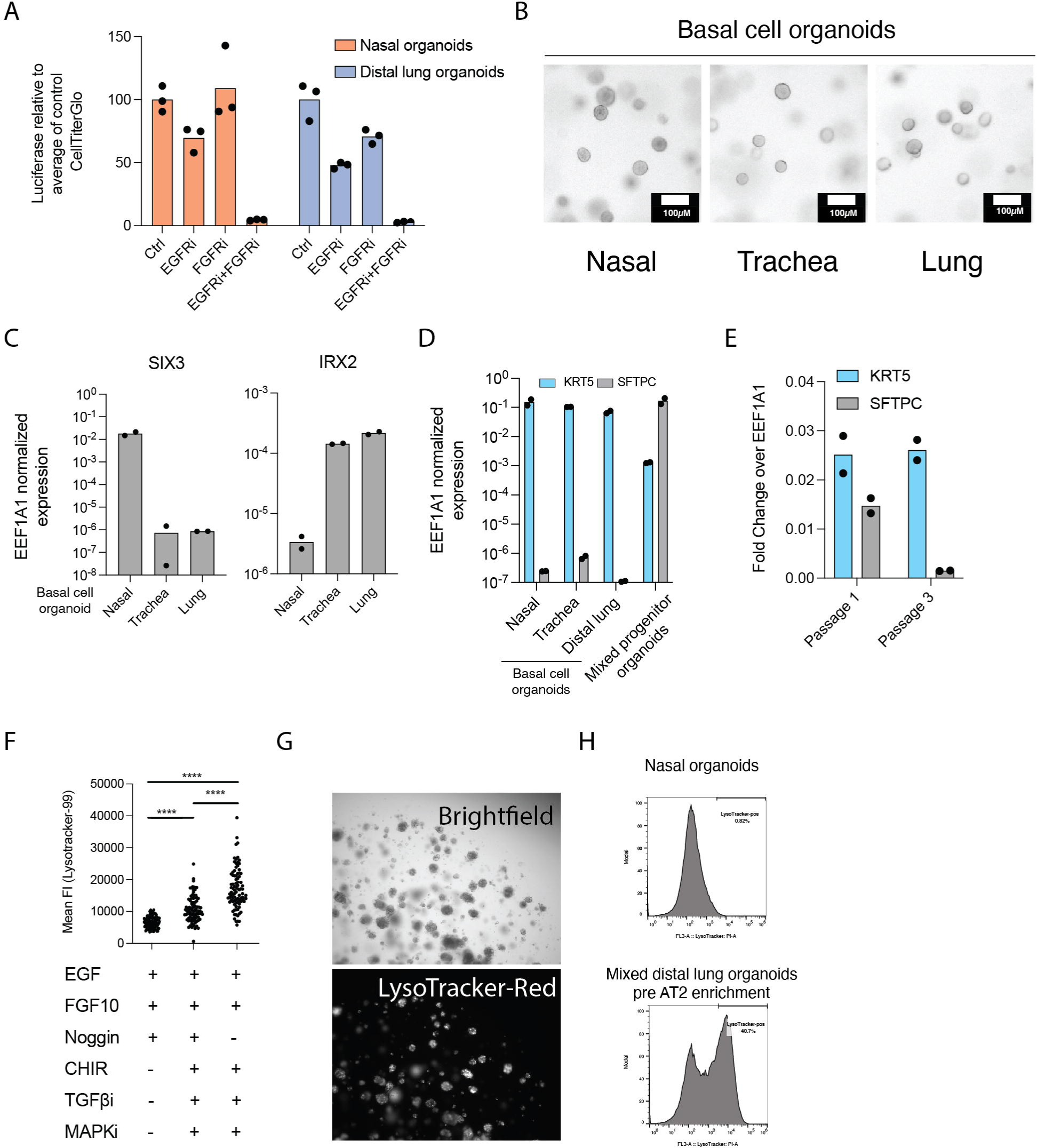
Establishment of bat airway organoids. A) Normalized CellTiter Glo luciferase viability assay for bat nasal or lung basal cell organoids grown in expansion medium with or without EGFR/FGFR inhibitors. Viability was normalized to the average luciferase of control (DMSO) treated organoids in nasal or lung cultures. Dots correspond to individual samples. B) Brightfield images of bat basal cell organoids grown from nasal, tracheal or distal lung tissue grown in basal cell expansion medium. C) RT-qPCR analysis of EEF1A1 reference gene normalized expression of SIX3 or IRX2 in bat basal cell airway organoids grown for six months. Reference gene normalized expression values are shown. Each dot represents a different organoid sample. D) RT-qPCR analysis of EEF1A1 reference gene normalized expression of KRT5 or SFTPC in bat basal cell organoids or mixed progenitor airway organoids. Reference gene normalized expression values are shown. Dots correspond to individual samples. E) RT-qPCR analysis of EEF1A1 reference gene normalized expression of KRT5 or SFTPC in bat distal lung organoids grown in complete distal lung organoid expansion medium and passaged three times at a ratio of 1:10 without FACS enrichment. Dots correspond to individual samples. F) Analysis of bat organoids stained with LysoTracker-Red dye prior to FACS sorting. Each dot represents the mean fluorescence intensity of one organoid obtained via Fiji Image analysis. The three groups represent different culture conditions. Unpaired Mann-Whitney tests were performed between conditions (**** P<0.0001). G) Fluorescence images of bat distal lung organoids cultured in complete distal lung organoid expansion medium and stained with LysoTracker-dye prior to FACS AT2 cell enrichment. H) FACS plots showing LysoTracker-Red fluorescent intensity distributions of single gated cells in bat nasal basal cell organoids grown in basal cell expansion medium, or distal lung organoids grown in complete distal lung organoid expansion medium. The percentages of LysoTracker-Red positive cells are shown.

**Supplemental Figure S2.**
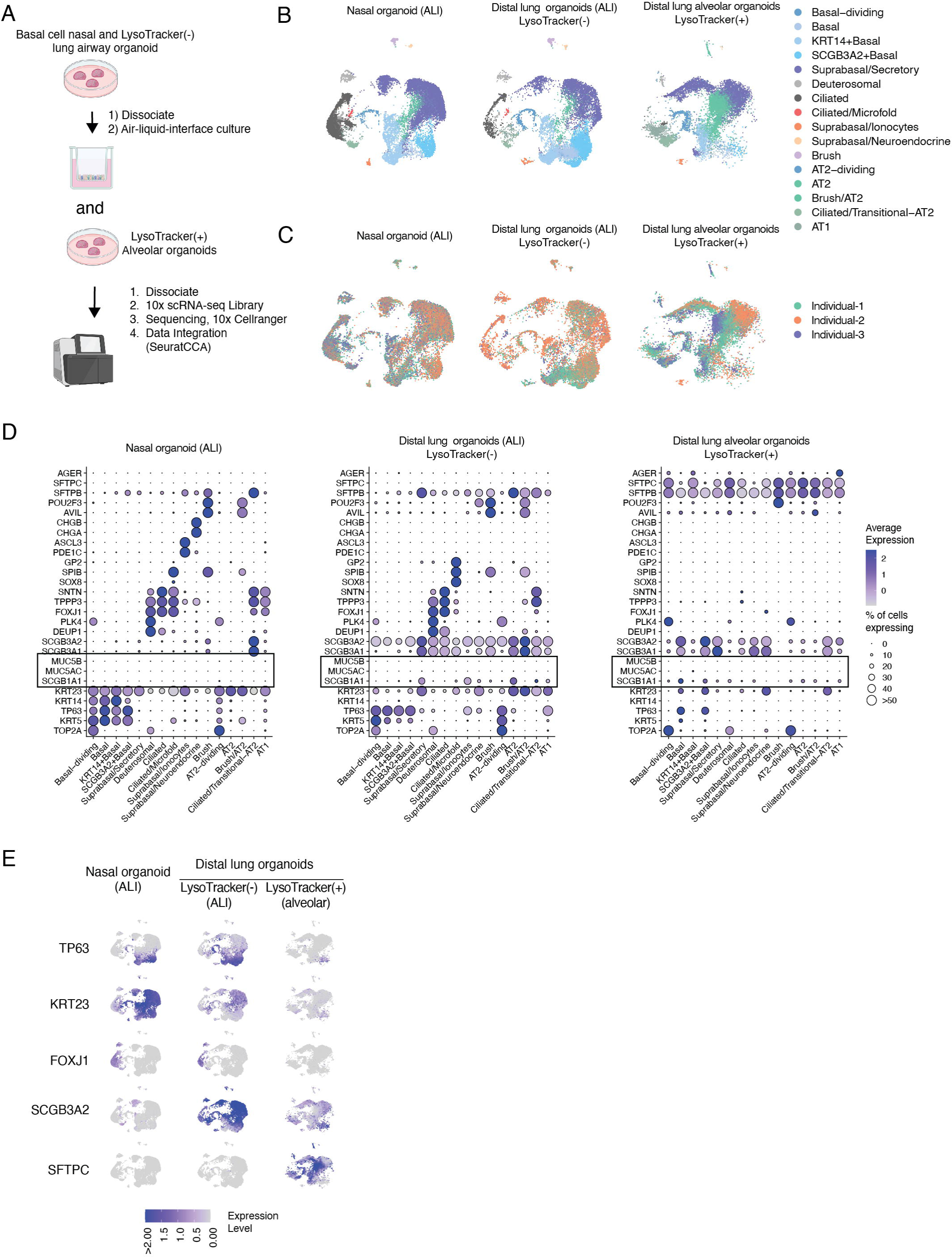
Single-cell RNA-sequencing of bat airway organoids. A) Schematic illustrating 10x single-cell RNA-sequencing experimental workflow. B) UMAP plot of integrated scRNA sequencing dataset showing cell type clusters split by the different organoid model identities. C) UMAP plot of integrated dataset showing cell type clusters split by different organoid models and colored by individual animal used for generating organoids. D) DotPlot analysis showing the average expression of markers for each cell cluster split by organoid identity. The dot sizes show the percentages of individual cell type expressing a given marker while the color intensities show the average expression values. Dot sizes were set to a maximum percentage of cells expressing the feature (dot size) of 50%. Genes expressed by more than 50% of cells have the same dot size. E) FeaturePlot showing the normalized expression of specific genes for each cell of an organoid model in the integrated UMAP space. The color indicates the expression level with a maximum normalized expression color cutoff set to 2. Different genes are shown in rows, different organoids models are arranged in columns and labeled above.

**Supplemental Figure S3.**
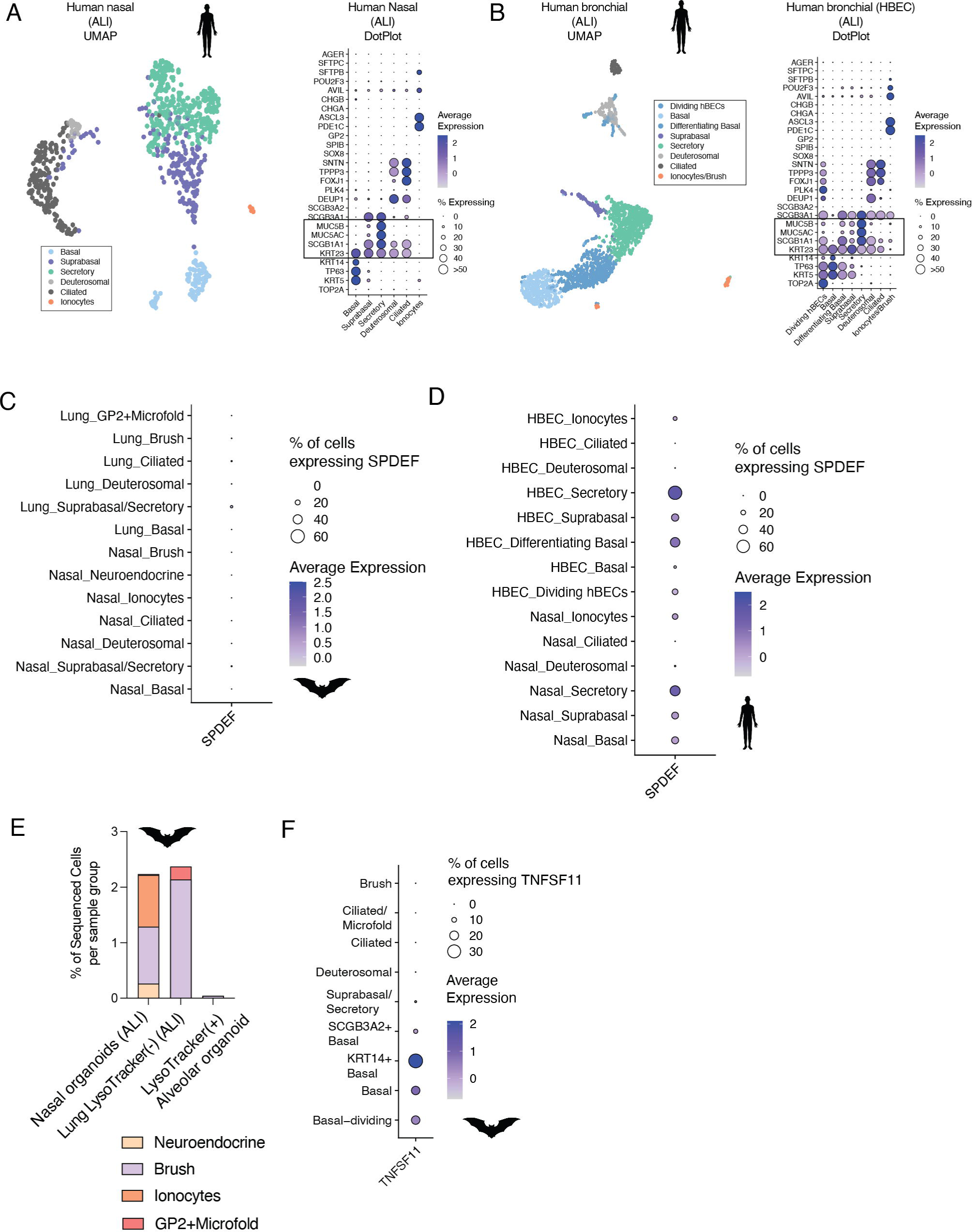
Single-cell RNA-sequencing of human airway cultures. A) Left: UMAP plot of single-cell RNA-sequencing data from human nasal air-liquid interface cultures. Right: DotPlot analysis showing the average expression of markers for each cell-cluster. The dot sizes show the percentages of individual cell type expressing a given marker while the color intensities show the average expression values. Dot sizes were set to a maximum percentage of cells expressing the feature (dot size) of 50%. Genes expressed by more than 50% of cells have the same dot size. B) Left: UMAP plot of single-cell sequencing data from human bronchial epithelium air-liquid interface cultures. Right: DotPlot analysis showing the average expression of markers for each cell-cluster. The dot sizes show the percentages of individual cell type expressing a given marker while the color intensities show the average expression values. Dot sizes were set to a maximum percentage of cells expressing the feature (dot size) of 50%. Genes expressed by more than 50% of cells have the same dot size. C) DotPlot analysis showing the average expression of SPDEF for each cell-cluster determined in bat organoids. The dot sizes show the percentages of individual cell type expressing a given marker while the color intensities show the average expression values. D) Same as in A) but for human organoids. E) Stacked barplot showing proportions of rare cell types found in each bat airway organoid model. F) DotPlot analysis showing the average expression of TNFSF11 (RANKL) in cell type of the bat distal lung organoid ALI cultures. The dot sizes show the percentages of individual cell type expressing a given marker while the color intensities show the average expression values.

**Supplemental Figure S4.**
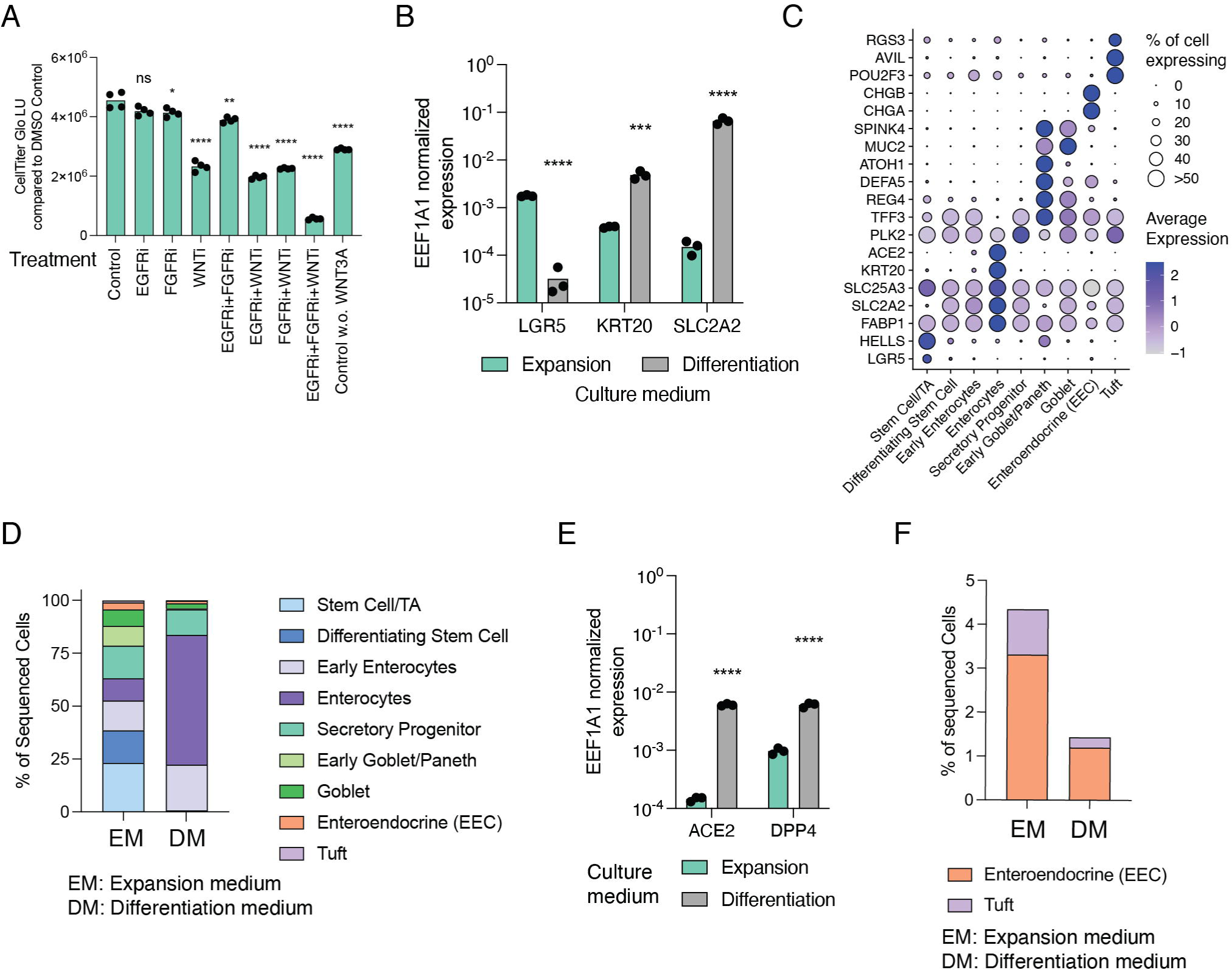
Bat Small Intestinal Organoid single-cell RNA-sequencing. A) CellTiter Glo Luciferase intensities for lysates of organoids cultured under different conditions. Unpaired student t-tests were performed between control and treatments in quadruplicate samples (ns: not significant, * P<0.05, ** P<0.01, *** P<0.001, **** P<0.0001). EGFRi (PD153035, 1µM),FGFRi (1µM Futibatinib), WNTi (1µM XAV-939). B) RT-qPCR analysis of EEF1A1 reference gene normalized expression of LGR5, KRT20 or SLC2A2 in bat small intestinal organoids cultured in expansion or differentiation medium. Unpaired student t-tests were performed between the two media conditions from triplicate samples (*** P<0.001, **** P<0.0001). C) DotPlot analysis showing the average expression of markers for each cell-cluster. The dot size shows the percentage of individual cell type expressing a given marker while the color intensity showing the average expression value. Dot sizes were set to a maximum percentage of cells expressing the feature (dot size) of 50%. Genes expressed by more than 50% of cells have the same dot size. D) Proportion of cell types found in single-cell RNA-sequencing data of bat small intestinal organoids culture in expansion (EM) or differentiation medium (DM). E) RT-qPCR analysis of reference gene normalized expression of ACE2 or DPP4 in bat small intestinal organoids cultured in expansion or differentiation medium. Unpaired student t-tests were performed between the two media conditions (**** P<0.0001). Each dot represents a different sample. F) Proportion of rare cell types found in single-cell RNA-sequencing data of organoids culture in expansion (EM) or differentiation medium (DM).

**Supplemental Figure S5.**
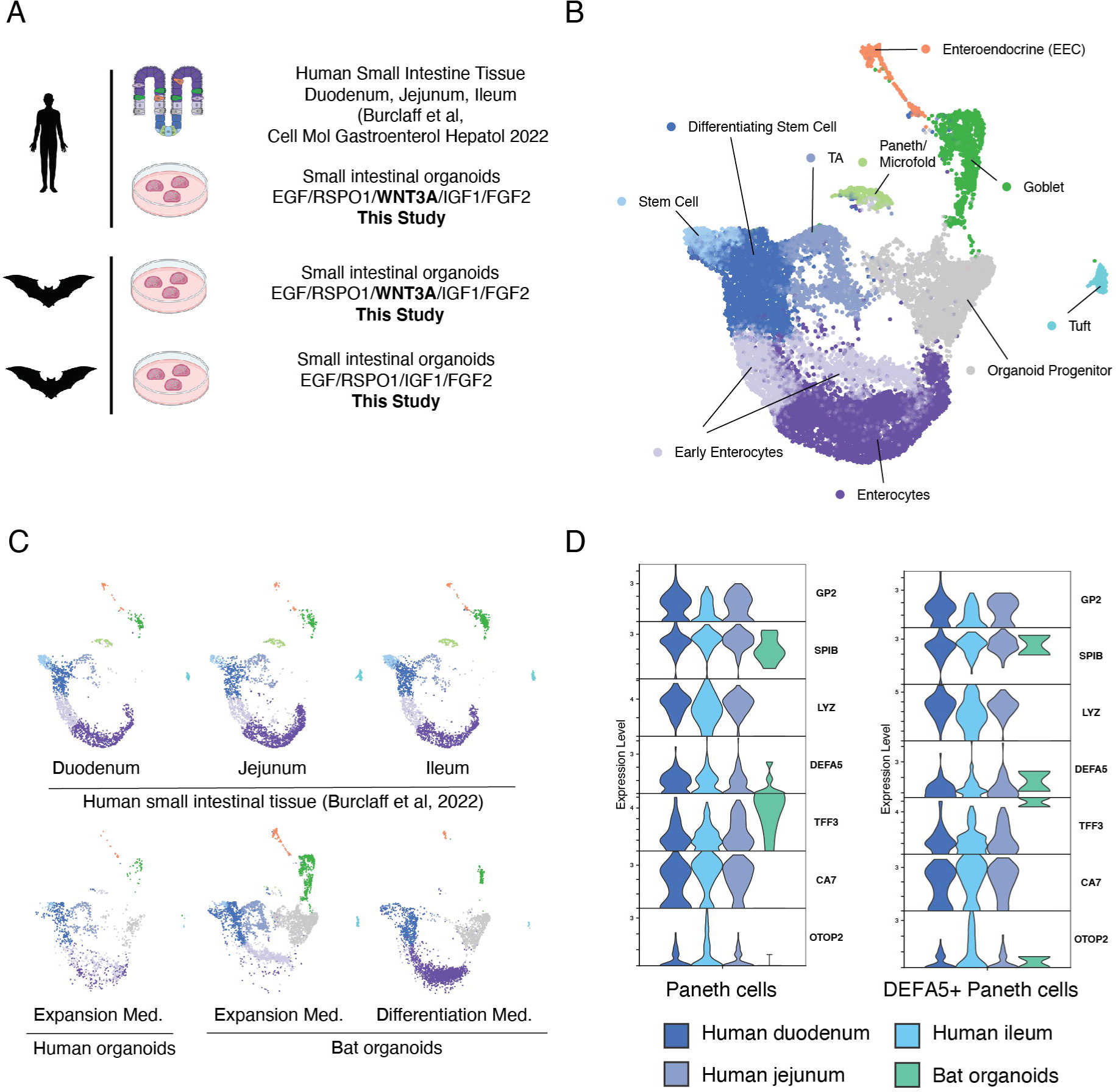
Intestinal human and bat organoid comparison. A) Schematic of cell types found in small intestines and which samples were used for single-cell RNA-sequencing data integration. Human small intestinal tissue data was derived from an external dataset (Burclaff et al., 2022)^68^. B) UMAP plot of integrated dataset showing assigned cell type clusters. C) UMAP plot of integrated dataset showing assigned cell type clusters, split by sample group identity. D) VlnPlot analysis of paneth cell marker (OTOP2, CA7, TFF3, DEFA5) or microfold cell marker (SPIB, GP2) are shown for mixed paneth/microfold cell cluster (left) and selected DEFA5+ paneth/microfold cells (right). The expression distribution is derived from individual cells.

**Supplemental Figure S6.**
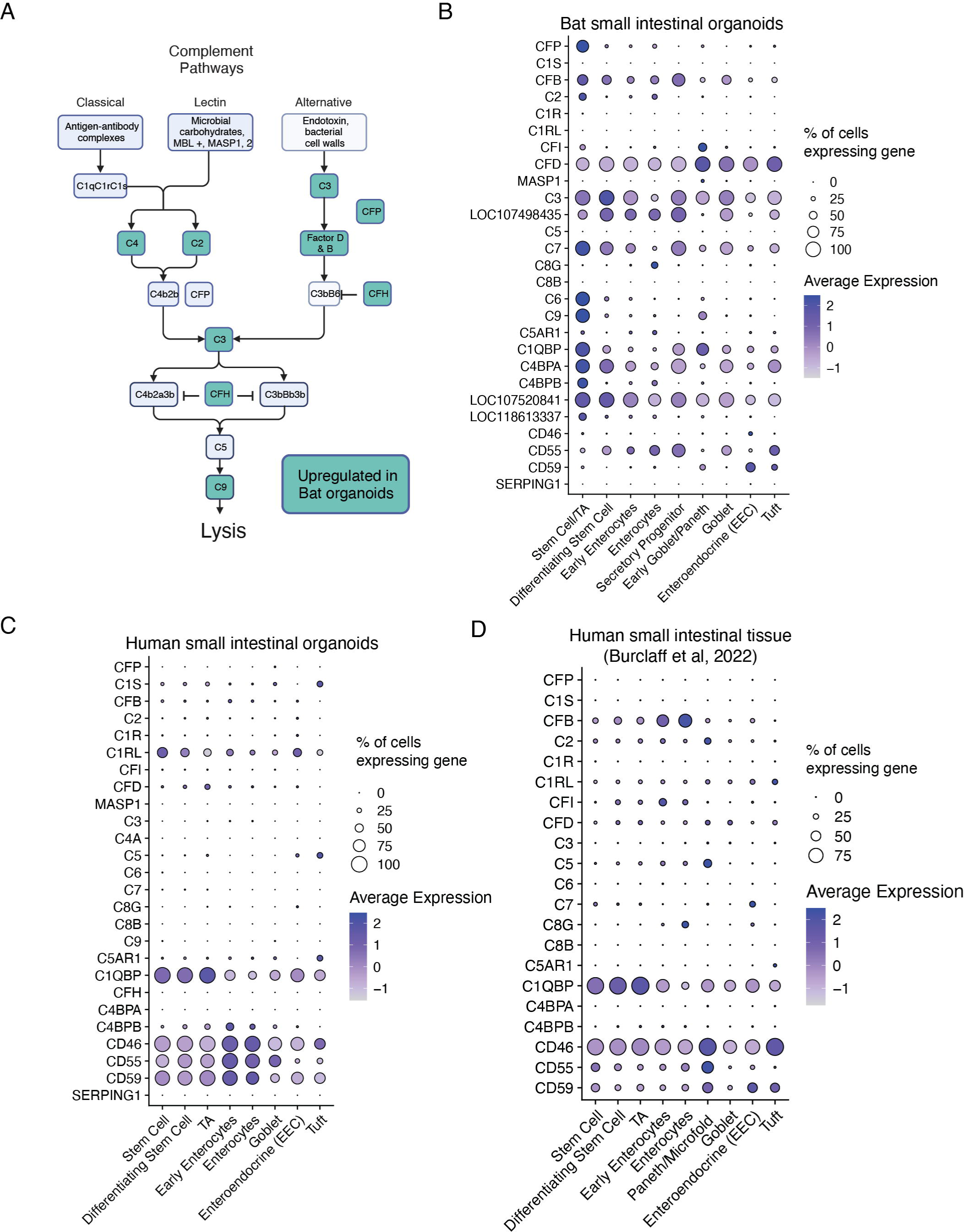
Expression of complement genes in bat and human small intestinal organoids and human small intestinal tissue. A) Schematic of cell genes involved in the classical or alternative complement system. Green boxes are genes upregulated in bat organoids. B) DotPlot analysis showing the average expression of complement genes for each cell cluster. The dot sizes show the percentages of individual cell types expressing a given marker while the color intensities show the average expression values. C) Same as in B) but for human small intestinal organoids. D) Same as in B) but for human small intestinal tissue.

**Supplemental Figure S7.**
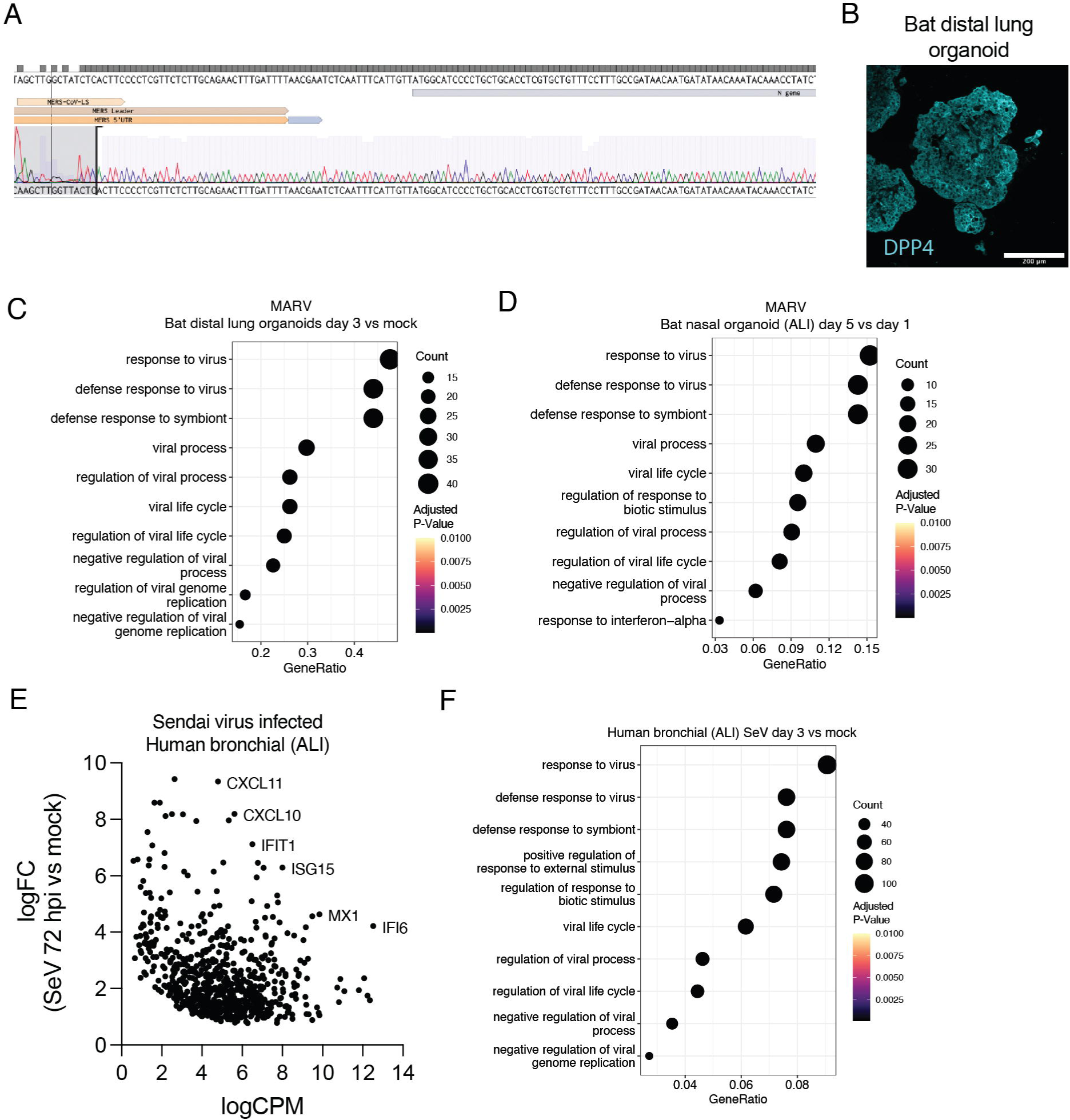
RNA virus infections in organoids. A) Sanger-sequencing result of the PCR product obtained with primers for the subgenomic transcript of the MERS Nucleocapsid RNA in infected bat small intestinal organoids. B) Immunofluorescence staining with anti-DPP4 antibody in bat distal lung alveolar organoids. C) Biological process enrichment analysis performed with *clusterProfiler* of differentially expressed genes in MARV infected versus uninfected bat distal lung organoids. The top 10 enrichment biological processes are shown. The values on the x-axis are the ratio of enriched to total genes for each biological process, while the dot size shows the number of enriched genes in the category. The color shows the adjusted P-value. D) Biological process enrichment analysis performed with clusterProfiler of differentially expressed genes in MARV infected versus uninfected bat nasal ALI cultures. The top 10 enrichment processes are shown. The values on the x-axis are the ratio of enriched to total genes for each biological process, while the dot size shows the number of enriched genes in the category. The color shows the adjusted P-value. E) Results from EdgeR differential gene expression analysis of human bronchial epithelial ALI cultures infected with Sendai virus (SeV) for 72 hours compared to uninfected samples. Each dot represents the expression and log-fold change (logFC) of an upregulated gene between infected and uninfected cultures. Positive logFC values mean upregulated in infected versus uninfected samples. Individual genes (ISGs) are highlighted. F) Biological process enrichment analysis performed with *clusterProfiler* of significantly upregulated genes in Sendai virus infected versus uninfected human bronchial epithelial ALI cultures. The top 10 enrichment processes are shown. The values on the x-axis are the ratio of enriched to total genes for each biological process, while the dot size shows the number of enriched genes in the category. The color shows the adjusted P-value.

**Supplemental Figure S8.**
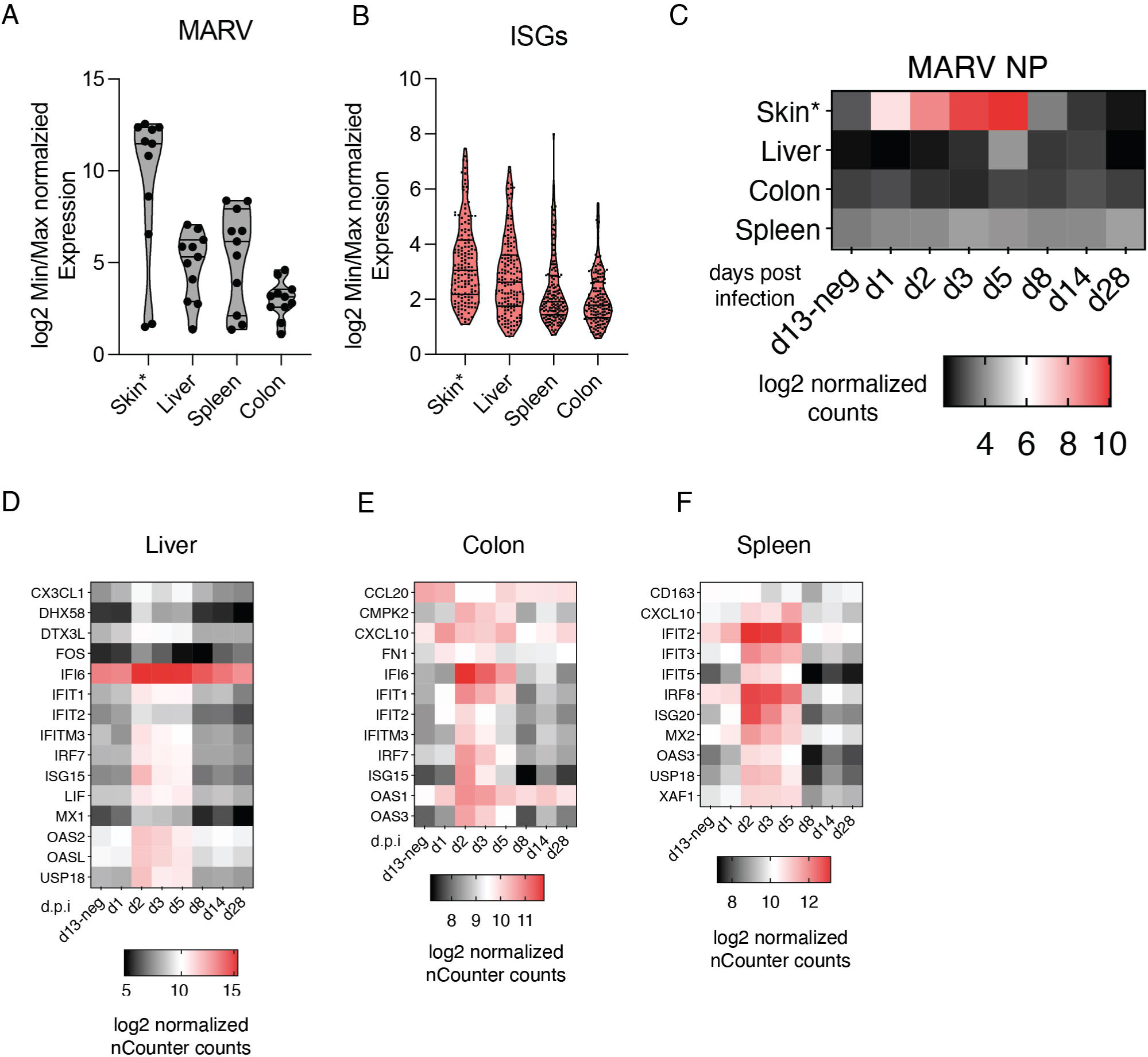
Re-analyses of *in vivo* MARV Egyptian fruit bat infections (Guito et al, 2021)^7^. A) Dynamic MARV expression in infected bats. Log2-transformed MARV specific nCounter CodeSet normalized expression changes of in infected bats throughput the infection time course. Normalizations were done by calculating the ratio between the maximum and minimum observed expression value. Each dots represent a different MARV gene measured. Each group represent a different tissue. B) Same as in A) but for interferon stimulated genes. C) Log2-transformed MARV-NP nCounter normalized expression in four different tissues throughout the infection time-course. D) Log2-transformed nCounter normalized expression of selected, dynamically regulated ISGs in liver tissue throughout the infection time-course. E) Log2-transformed nCounter normalized expression of selected, dynamically regulated ISGs in colon tissue throughout the infection time-course. F) Log2-transformed nCounter normalized expression of selected, dynamically regulated ISGs in spleen tissue throughout the infection time-course.

**Supplemental Figure S9.**
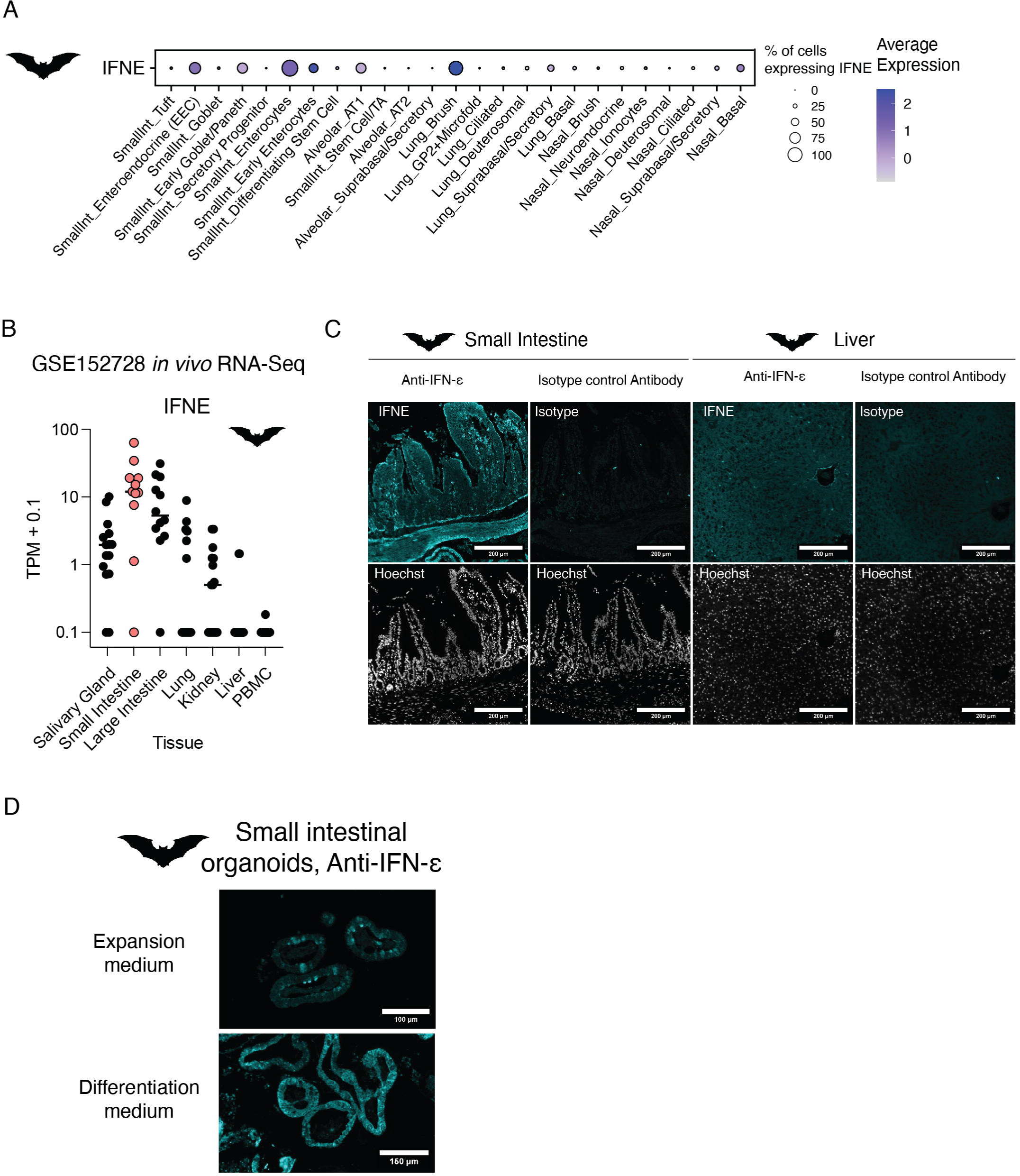
Interferon-ε expression in Egyptian fruit bat organoids. A) DotPlot expression analysis showing the average expression of interferon genes for each bat organoid cell type determined by single-cell RNA-sequencing. The dot size shows the percentage of cells in each cell type expressing IFNE. B) Dotplot showing TPM+0.1 normalzied RNA expression values for IFNE in different tissue from external bulk RNA-sequencing data (GSE152728^88^). Each dot represents an individual Egyptian fruit bat tissue sample. C) Immunofluorescence staining with anti-Interferon-ε antibody or isotype-control antibody in Egyptian fruit bat small intestine or liver tissue. DAPI counterstaining is shown below. D) Immuno-fluorescence staining with anti-Interferon-ε antibody in bat small intestinal organoids grown in expansion (top) or differentiation (bottom) medium.

**Supplemental Figure S10.**
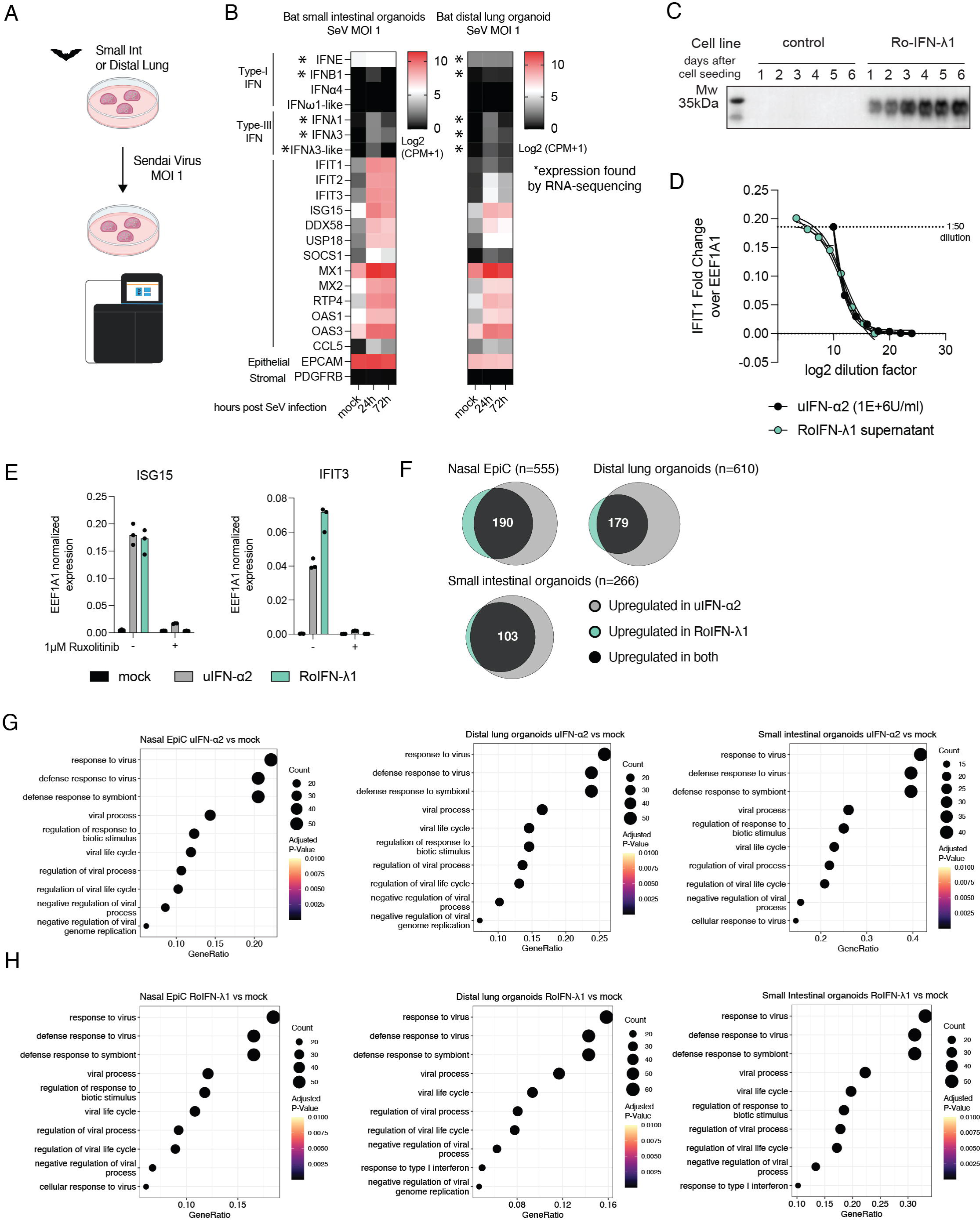
Interferon stimulation of bat organoids. A) Schematic of experimental overview of Sendai virus infection in bat organoids. B) Heatmap showing the average log2 normalized RNA expression (in log2 EdgeR CPM) for interferon and interferon stimulated genes in two different organoid models after Sendai virus infection. Each value presents the average of three replicates at different time-points post infection and mock infected. C) Western-blot analysis of culture supernatant from HEK293T cells stably expressing His-tagged RoIFN-λ1. Control supernatant is from unmodified HEK293T cells. Results are shown for culture supernatants from different days after seeding cells (see methods). D) Dose-response curve of IFIT1 expression measured by RT-qPCR in primary bat nasal cells stimulated with different amounts of universal IFNα2 or RoIFN-λ1 supernatant harvested on day 3 post seeding. EEF1A1 reference gene normalized IFIT1 expression is shown. A curve was fitted in Graphpad prism. 1:50 dilution of RoIFN-λ1supernatant was determined to yield equivalent activation of IFIT1 compared to 1000 U/ml universal Interferon-α2. E) RT-qPCR analysis of EEF1A1 reference gene normalized expression of ISG15 and IFIT3 in primary bat nasal cells stimulated for eight hours with 1000 U/ml universal IFN-α2 or equivalent functional amounts of RoIFN-λ1 supernatant in the presence or absence of 1 µM Ruxolitinib. Each dot represents a biological replicate. F) Venn diagrams showing the results from differential gene expression analysis comparing uIFN-α2 and RoIFN-λ1 stimulations in bat organoids. The total number of interferon upregulated genes is labeled with the sample type on the top. A common set of upregulated genes is highlighted by the black area formed by the intersection of both treatments for each sample. The numbers of commonly upregulated genes are shown. G) Biological process enrichment analysis performed with *clusterProfiler* of significantly upregulated genes in IFN-α2 treated versus untreated cells. The top 10 enriched bioloigical processes are shown. The values on the x-axis are the ratio of enriched to total genes in each bioloigical process, while the dot size shows the number of enriched genes. The color shows the adjusted P-value. The tested conditions are labeled on top. H) Same as for F) but for RoIFN-λ1 treated cells and organoids.

**Supplemental Figure S11.**
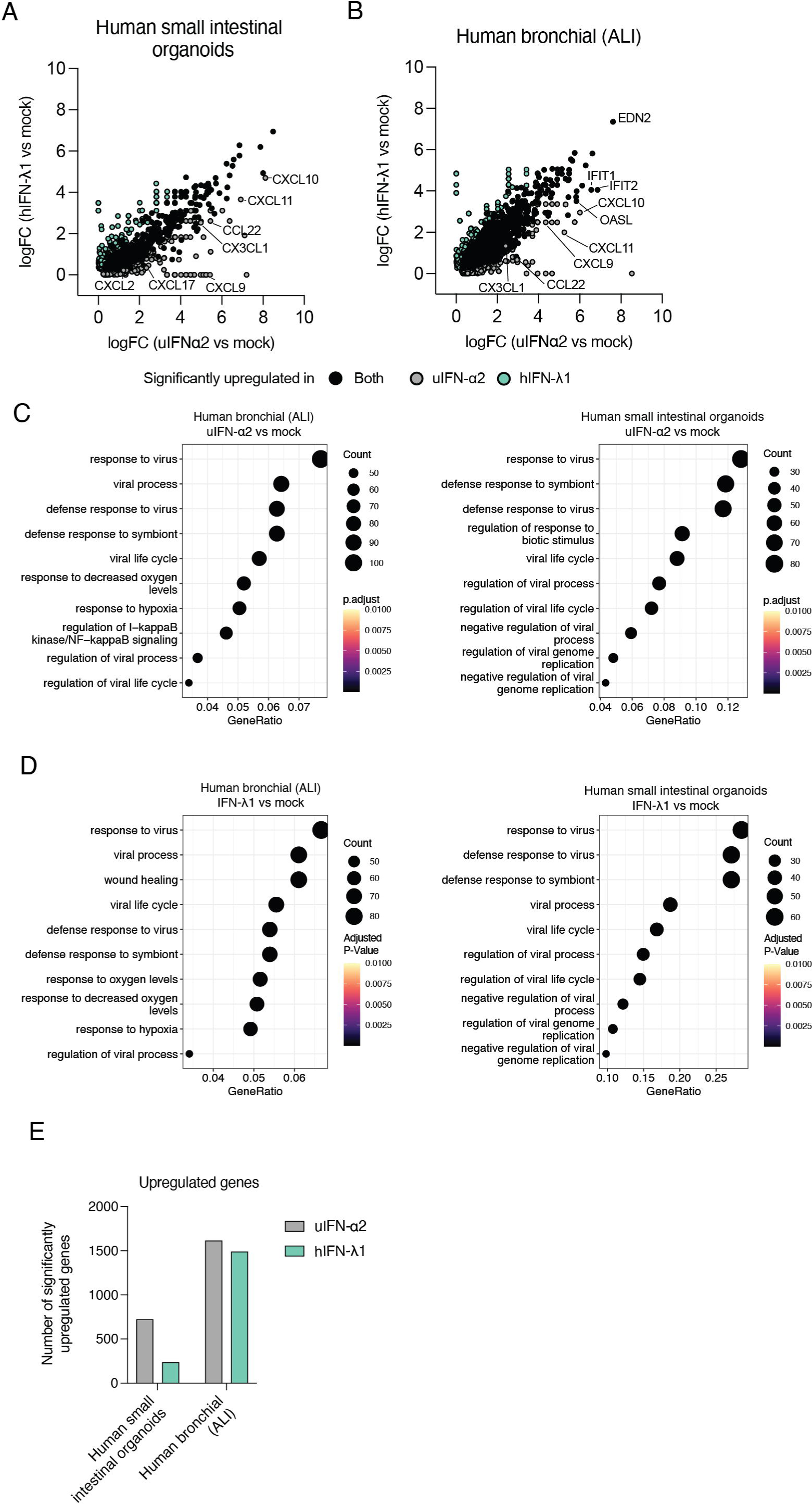
Interferon stimulation studies in human organoids. A) Scatterplot showing the log-fold change of genes determined by comparing the expression of interferon to mock treated human small intestinal organoids. Each dot represents a pairwise-comparison in either IFNα2 to mock or hIFN-λ1 to mock treated cells. Genes significantly upregulated after both uIFN-α2 and IFN-λ1 interferon stimulation or only by one are colorcoded. Individual genes are highlighted. B) Same as for A) but for human bronchial ALI cultures. C) Biological process enrichment analysis performed with clusterProfiler of significantly upregulated genes in IFN-α2 treated versus untreated cells. The top 10 enriched bioloigical processes are shown. The values on the x-axis are the ratio of enriched to total genes in each bioloigical process, while the dot size shows the number of enriched genes. The color shows the adjusted P-value. The tested conditions are labeled on top.The tested conditions are labeled on top. D) Same as in C) but for IFN-λ1 treatments. E) Barplot showing the number of significantly upregulated genes in human organoid culture systems following interferon-α2 or hIFN-λ1 treatment.

**Supplemental Figure S12.**
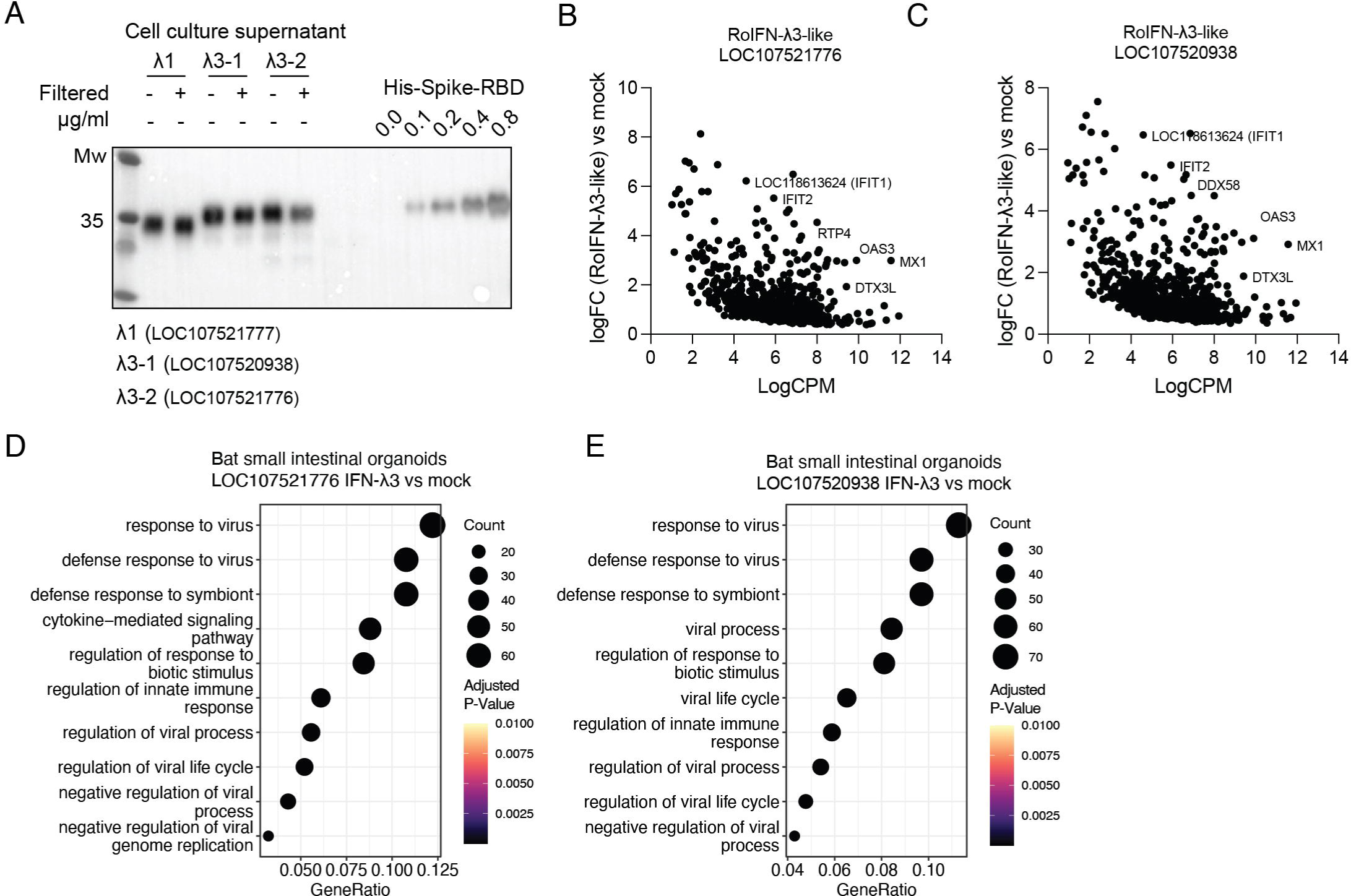
RoIFN-λ3-like stimulations in bat organoids. A) Western-blot analysis of culture supernatant from HEK293T cells stably expressing His-tagged RoIFN-λ-like genes. The filtered or unfiltered supernatant as well as a dilution series of his-tagged SARS-CoV-2 Spike-RBD protein are shown. Different expressed RoIFN-λ-like genes are labeled with the respective NCBI gene ID. B) Results from differential gene expression analysis of bat small intestinal organoids treated with LOC107521776 derived supernatant compared to mock treated conditions. Each dot represents the expression and log-fold change of an upregulated gene between IFN treated or mock treated samples. Positive LogFC values means upregulated in IFN treated organoids. Individual ISGs are highlighted. C) Same as in B) but for IFN-λ3-like gene LOC107520938 D) Related to experiment shown in B). Biological process enrichment analysis performed with *clusterProfiler* of significantly upregulated genes in LOC107521776 treated versus untreated cells. The top 10 enrichment processes are shown. The values on the x-axis are the ratio of enriched to total genes in each group, while the dot size shows the number of enriched genes. The color shows the adjusted P-value. The tested conditions are labeled on top. E) Related to experiment shown in C). Same as in D) but for IFN-λ3-like gene LOC107520938.

**Supplemental Figure S13.**
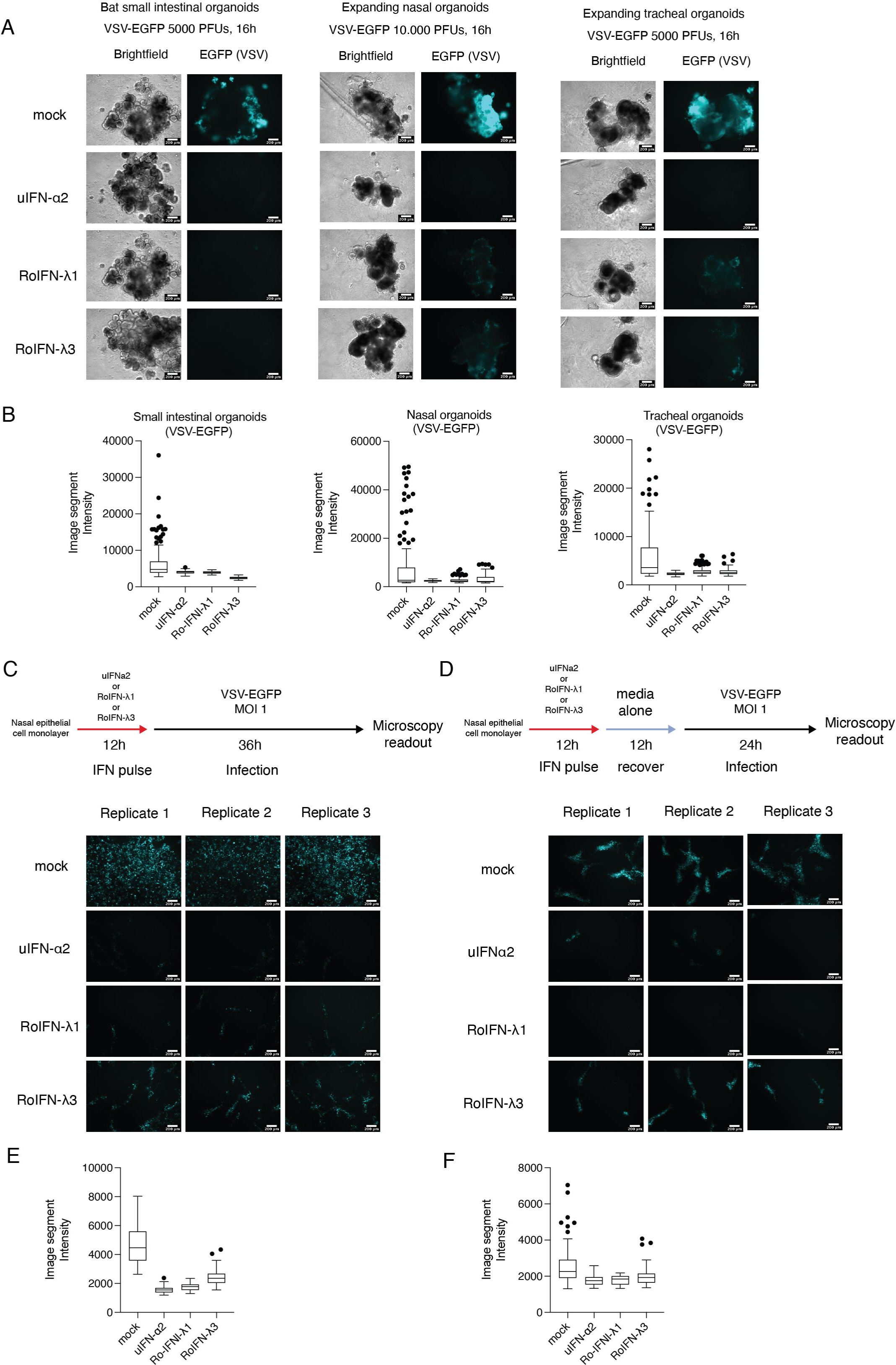
Interferon-mediated antiviral immunity in bat organoids. A) Brightfield and epifluorescence images of infected bat small intestinal, nasal or tracheal organoids with VSV-EGFP. Organoids were stimulated with universal interferon-α2, RoIFN-λ1, RoIFN-λ3 or mock treated, starting 12 hours before VSV-EGFP infection. Organoids were imaged 16 hours post infection. B) ImageJ quantification of organoids shown in A). EGFP signal intensities of individual image segments are shown for each condition and shown as Tukey boxplots with outlier outside the 1.5 interquartile range (IQR) highlighted (see methods). C) Antiviral protection of nasal organoid derived monolayer cultures pre-treated for 12 hours with universal interferon-α2, RoIFN-λ1, RoIFN-λ3 or mock treated before infection with VSV-EGFP at an MOI of 1 (experimental workflow found on top). Epifluorescence images are shown for cells infected for 36 hours in triplicates. D) Antiviral protection of nasal organoid derived monolayer cultures treated for 12 hours with universal interferon-α2, RoIFN-λ1, RoIFN-λ3 or mock treated. Cells were then incubated in media not containing interferons for an additional period of 12h before infection with VSV-EGFP at an MOI of 1 (experimental workflow found on top). Epifluorescence images are shown for cells infected after 24 hours in triplicates. E) ImageJ quantification of cells shown in C). EGFP signal intensities of individual image segments are shown for each condition and shown as Tukey boxplots with outlier outside the 1.5 interquartile range (IQR) highlighted (see methods). F) ImageJ quantification of cells shown in D). EGFP signal intensities of individual image segments are shown for each condition and shown as Tukey boxplots with outlier outside the 1.5 interquartile range (IQR) highlighted (see methods).

**Supplemental Figure S14.**
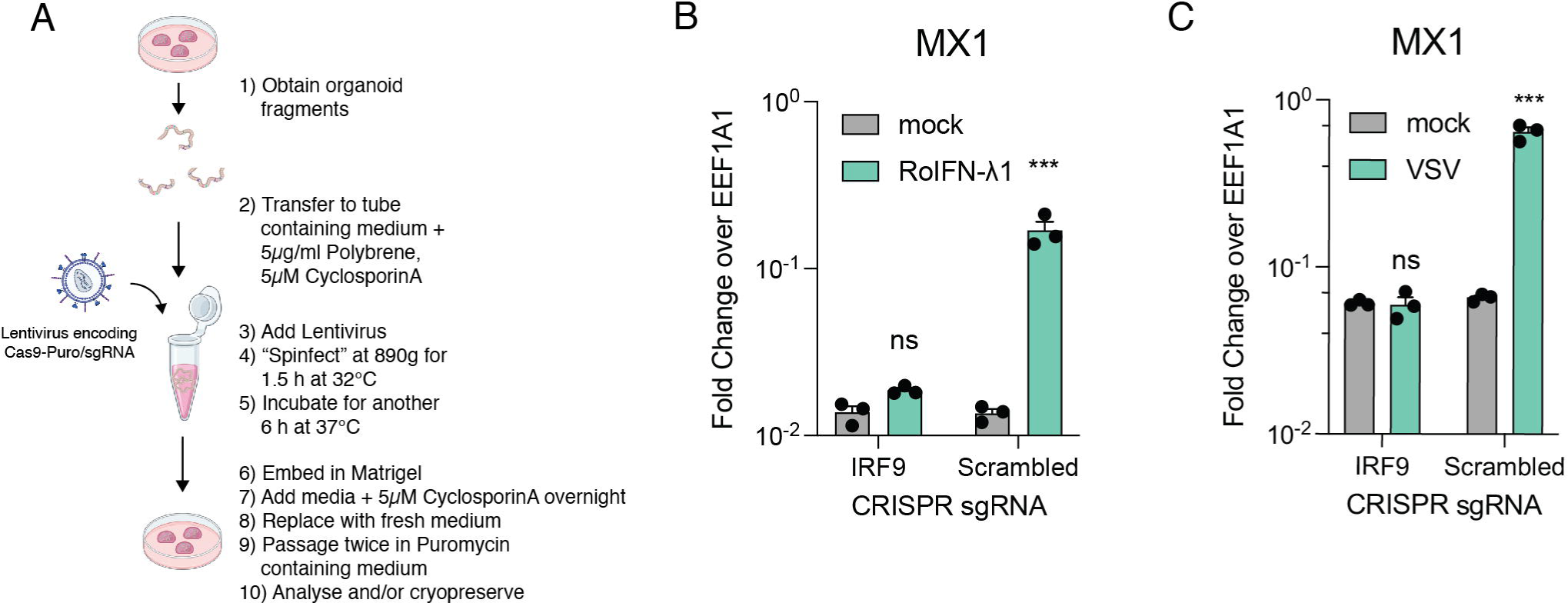
CRISPR-Cas9 in bat organoids. A) Schematic of CRISPR-Cas9 workflow in bat organoids. B) RT-qPCR expression analysis of MX1 (an ISG) in bat small intestinal organoids expressing Cas9 and a scrambled guide RNAs treated IFN-λ1 for eight hours. Reference gene normalized expression values for MX1 are shown. Each dot represents a biological replicate. Unpaired student t-tests were performed between mock and treatments (ns: not significant, * P<0.05, ** P<0.01, *** P<0.001, **** P<0.0001). C) RT-qPCR expression analysis of MX1 in bat small intestinal organoids expressing Cas9 and a scrambled guide RNA infected with VSV-EGFP at an MOI of 0.05 for 72 hours. Reference gene normalized expression values for MX1 are shown. Each dot represents a biological replicate. Unpaired student t-tests were performed between mock and infections (ns: not significant, * P<0.05, ** P<0.01, *** P<0.001, **** P<0.0001).

## Materials and Methods

### Bat and human and tissue and cell collection

*Rousettus aegyptiacus* bat tissue for organoid generation was derived from a breeding colony at the Friedrich-Loeffler-Institute in Germany. Sampling was performed in accordance with current european and national animal welfare regulations, after ethical review and approval by the authority of the federal state of Mecklenburg-Western Pomerania, Germany, and the experiments were carried out according to ARRIVE guidelines (https://arriveguidelines.org). Freshly harvested organ tissue from male and female bats was washed in PBS and cut into 3-5 mm small pieces before placing up to four individual pieces into a 1 ml cryostorage tube containing 0.5ml-1ml of Cryostore10 freezing medium (Stemcell Technologies). Vials were immediately transferred to a cryostorage container (Mr. Frosty) and frozen overnight at –70°C, before long-term liquid nitrogen storage. In total, nasal respiratory, lung and small intestinal tissue from three individual bats, and tracheal tissue from two individual bats were obtained and used in this study.

Human nasal epithelial cells were collected from a brush biopsy of the mid-turbinate section of a healthy female donor and placed into Advanced DMEM/F12 (Gibco). After collection of cells via centrifugation, individual cells were obtained by TrypLE express (Gibco) dissociation, followed by plating onto PureCol (Advanced BioMatrix, diluted 1:30 in PBS) treated cell culture dishes in PneumaCult-Ex plus expansion medium (Stemcell Technologies). Nasal epithelial cells were expanded for up to two additional passages in PneumaCult-Ex plus expansion medium and cryopreserved in liquid nitrogen in Cryostore10 freezing medium. Experiments were approved by the ethic commission of the Medical University of Vienna (Votum ECS 2234/2021). Air-liquid-interface quality controlled human bronchial epithelial cells were purchased from Promocell (PC-C-12640, Lot 446Z036.9) and were initially recovered in PromoCell airway expansion medium. Cells were further expanded for up to two additional passages in PromoCell airway expansion medium or PneumaCult-Ex plus expansion medium and cryopreserverd in Cryostore10 freezing medium.

### Establishment and maintenance of bat airway organoids

Frozen tissue pieces were quickly thawed in a water bath before transferring into a GentleMACS dissociation tube (Miltenyi Biotec) containing Advanced DMEM/F12 + 1:100 Glutamax (Gibco) + 10 mM HEPES (Gibco) + DNaseI (Roche) + 1.25 µg/ml collagenase from *Clostridium histolyticum* (Sigma). A homogenous cell suspension was obtained by initial disruption of bulk tissue pieces using program m_lung_01, followed by dissociation using protocol 37C_mLIDK_01 in a GentleMACS dissociator (Miltenyi). Dissociation was stopped by addition of 10 % FBS (Gibco) in Advanced DMEM/F12. The cell suspension was strained through a 70 µm filter and cells were collected by centrifugation. To remove red blood cells, the cell pellet was re-suspended and incubated in 1 ml of Red Blood Cell Lysis Buffer (Roche) for 5 minutes. Cells wereresuspended in Advanced DMEM/F12 and counted using Trypan blue to estimate live/dead cell number percentages. If the cell viability was below 60 %, dead cells were removed by AnnexinV-based magnetic bead selection following the manufactureŕs instructions (Stemcell Technologies). In short, cells were collected by centrifugation, resuspended in PBS+2 % FBS (Gibco) + 1 mM CaCl2 (Sigma) and transferred to a 5 ml FACS tube. 50 µl of AnnexinV-antibody and Biotin-beads were added and incubated for 5 minutes. 50µl of RapidSphere were added for 3 minutes. 1.5 ml of PBS + 2% FBS (Gibco) + 1 mM CaCl2 (Sigma) was added and the tube subsequently placed into a magnet. After 3-minute separation, unbound viable cells were decanted, collected by centrifugation, and resuspended in Advanced DMEM/F12 before counting.

For nasal and tracheal epithelial tissues, cells were either expanded in 2D using PneumaCult-Ex plus expansion medium as described for human airway epithelial cells (see above) or expanded in 3D culture conditions. For 3D culture establishment, 25.000 viable cells were resuspended in 50 µl of ice-cold matrigel for organoid culture (Corning, Cat. # 356255) and platted onto pre-warmed 24-well plate cell culture dishes. After solidification, basal cell expansion medium containing Advanced DMEM/F12 + 1:100 Glutamax (Gibco) + 10 mM HEPES (Gibco) + 1.25 mM N-Acetyl-Cystein (Sigma) + 1:50 B-27 Supplement (Gibco) + 1:100 N-2 Supplement (Gibco) + 50 ng/ml human EGF (Gibco) + 25 ng/ml human FGF10 (Stemcell Technologies) + 25 ng/ml human Noggin (Stemcell Technologies) + 10 vol% conditioned medium containing Rspondin1 (produced from HA-R-Spondin1-Fc HEK293T cell line, Trevigen) + 500 nM A-8301 (Selleckchem) + 1:500 Primocin (Invivogen) + 10 µM Y-27632 ROCK inhibitor (Stemcell Technologies) (only for the first two days) was added. Media was changed every 2-3 days. Organoids were split by dissociation of cells and re-seeding 5000 cells per 50 µl matrigel droplet every 7-9 days. Expanding organoids were then used for differentiation or cryopreserved using Cryostore10 freezing medium. For differentiation, 100.000 nasal epithelial cells were seeded in basal cell expansion medium or PneumaCult-Ex plus expansion medium + 10µM Y-27632 onto a PureCol treated 24-well plate size, 0.4 µm pore size Transwell insert (Corning). Apical and basal chamber medium was replaced every 2-3 days until cells reached confluency, after which they were air-lifted by removing medium in the apical chamber and replacing the basal chamber medium with complete PneumaCult air-liquid interface medium (Stemcell Technologies) + 1:500 Plasmocin. Cultures were differentiated at the air-liquid interface for 30-40 days with media changes every 3-4 days, and weekly apical PBS washes to remove mucus and/or dead cells. Alternatively, organoids were differentiated in 3D by replacing the expansion medium with PneumaCult air-liquid interface medium + 10 µM DAPT (Sigma) 4-5 days after initial expansion, for a period of 15-20 days. To prevent loss of differentiating 3D organoids due to attachment to the plastic dish, organoids were collected in Cell Recovery Solution (Corning) and left on ice for 30 minutes to remove matrigel. Organoids were then re-seeded into fresh matrigel and overlayed with PneumaCult air-liquid interface medium + 10 µM DAPT. This step was necessary after a total incubation period of ∼10 days. To determine growth factor signaling requirements, inhibitors of EGFR (PD153035, 1 µM) and/or FGFR (1 µM Futibatinib) were added to the complete culture medium after passaging. Media containing inhibitors was changed every 2 days. After 7 days, organoids were collected and lysed in 100 µl of a 1:1 dilution containing CellTiterGlo (Promega) and PBS. Luciferase was read-out after 5-minute incubation on a microplate reader with 500 ms exposure.

Distal lung 3D organoids were established by seeding 25.000 dissociated viable distal lung cells in 50 µl matrigel droplets, which after solidification were overlayed by basal medium containing Advanced DMEM/F12 + 1:100 Glutamax + 10 mM HEPES + 1.25 mM N-AcetylCystein + 1:50 B-27 Supplement + 1:100 N-2 Supplement + 50 ng/ml human EGF + 1:500 Primocin + 0.002% Heparin (Sigma), supplemented with additional factors as described in the main text. For a complete distal lung expansion medium containing both basal stem cell and AT2-cells, 25 ng/ml human FGF10 (Stemcell Technologies) + 25 ng/ml human Noggin (Stemcell Technologies) + 3 µM CHIR9902 (Stemcell Technologies) + 10 µM SB-431542 (Stemcell Technologies) + 1 µM BIRB796 (Stemcell Technologies) were added to the culture medium. Cultures were supplemented with 10 µM Y-27632 for the first three days after culture establishment and passaging. Organoids were expanded for one additional passage, before separation of AT2+ and AT2-negative cells using LysoTracker™ Red DND-99-dye (Thermofisher) based sorting. To accomplish this, 1:10.000 diluted LysoTracker dye (Thermofisher) was added to the organoid cultures for 2 hours, followed by single cell dissociation and sorting of LysoTracker positive (AT2 enriched) and LysoTracker negative (basal cell enriched) lung epithelial progenitor cells. Sorted cells were placed in basal cell expansion medium (LysoTracker negative cells) or complete distal lung expansion medium (LysoTracker positive), supplemented with 10 µM Y-27632 for the first 3-days. Organoids were split by dissociation and re-seeded at 5000 cells per 50 µl matrigel droplet into new 24-well cell culture dishes. Distal lung basal cells were differentiated as described above for the nasal epithelial cells.

### Establishment and maintenance of bat small intestinal organoids

Bat small intestinal cells were obtained from frozen tissue as described for airway cells above. For the organoid establishment, 25.000 cells were seeded in 50 µl of matrigel and overlayed with small intestinal organoid expansion medium containing Advanced DMEM/F12 + 10 vol% conditioned medium containing Rspondin1 + 50 vol% conditioned medium containing human WNT3A (the L-Wnt3a cell line was a gift from Hans Clevers) + 1:100 Glutamax + 10 mM HEPES + 1 mM N-AcetylCystein + 1:50 B-27 Supplement + 50 ng/ml human EGF + 100 ng/ml human Noggin +100 ng/ml human IGF1 (Stemcell Technologies) + 50 ng/ml human FGF2 (Stemcell Technologies) + 500 nM A-83-01 + 1:500 Primocin + 10 µM Y-27632 (only for the first two days). Media was changed every 2-3 days. Passaging was done by collecting organoids grown in matrigel droplets in cold Advanced DMEM/F12 and quick centrifugation (short acceleration to 5000 rcf, ∼3-4 seconds). The supernatant containing dead cells and matrigel was removed, and 500 µl of TrypLE Express was added to resuspend the organoid pellet. After 3 minutes at room temperature, 500 µl of cold Advanced DMEM/F12 was added to the suspension and organoids were fragmented by manual pipetting. After centrifugal collection of organoid fragments, the supernatant was removed and 1000 µl of Advanced DMEM/F12 added. The organoid fragment suspensions were transferred to a new tube and cells collected by centrifugation. Finally, organoid fragments were seeded in ice-cold matrigel. We typically split small intestinal organoids once a week at a 1:6-1:12 split ratio.

To determine growth factor signaling requirements, inhibitors of EGFR (PD153035, 1 µM), FGFR (1 µM Futibatinib), WNT (1 µM XAV-939) were added to the complete culture medium after passaging. In parallel, a complete medium lacking WNT3A was used. Medium was changed every 2 days. After 7 days, organoids were collected and lysed in 100 µl of 1:1 dilution containing CellTiterGlo and PBS. Luciferase was read-out after 5-minute incubation on a microplate reader with 500ms exposure. Organoid differentiation was initiated by replacing the culture medium 3-4 days post passaging with differentiation medium containing Advanced DMEM/F12 + 10 vol% conditioned medium containing Rspondin1 + 1:100 Glutamax (Gibco) + 10 mM HEPES (Gibco) + 1 mM N-AcetylCystein (Sigma) + 1:50 B-27 Supplement (Gibco) + 50 ng/ml human EGF +100 ng/ml human IGF1 (Stemcell Technologies) + 50 ng/ml human FGF2 (Stemcell Technologies) + 500 nM A-8301 (Selleckchem) + 1:500 Primocin (Invivogen). Differentiation was carried out for addition 3-5 days before using organoids for experiments. Human small intestinal organoids were established from healthy duodenum biopsy specimens by isolating intestinal crypts (full ethical approval (REC-12/EE/0482)). We received a vial of frozen organoids^110^ and cultured them alongside bat organoids in expansion medium as detailed above.

### Production of Egyptian fruit bat interferon-λ

The coding sequences lacking the signal peptide of *Rousettus aegyptiacus* IFNL1-like (LOC107521777) and IFNL3-like (LOC107521776 or LOC107520938) were obtained by reverse-transcription and PCR from RNA of poly(I:C) (Invivogen) stimulated bat nasal epithelial cells. Stimulation was carried out by transfecting 30.000 cells with 100 ng of HMW poly(I:C) (Invivogen) in 10 µl Opti-MEM I (Gibco) + 0.15 µl Lipofectamine 3000 (Thermofisher) for 8 hours before harvesting of RNA. The PCR fragments were cloned into a PiggyBac mammalian expression vector (System Biosciences, PB210PA-1), modified to harbor a Secrecon-AA signal peptide^111^ for secretion at the N-terminus, and 6x-His-tag at the C-terminus. The expression vector was further modified to contain an IRES-Puromycin selection cassette. A stable IFN-λ-expressing cell line was obtained by transfection of one million Lenti-X (HEK293T, Takara) cells with 500 ng Super PiggyBac Transposase vector and 1250 ng of PiggyBac transposase mammalian expression vector using Lipofectamine 3000 following to the manufactureŕs instructions. Stable cells were selected with 1 µg/ml Puromycin and expanded for 2-weeks. For harvesting of RoIFN-λ, cells were seeded at high density (1.2 million cells per 1-well of a 6-well plate) and incubated for 3-days. The supernatant was cleared by centrifugation, filtered through a 0.22 µm syringe filter and stored in aliquots at –70°C. Biological activity was determined by adding diluted amounts of RoIFN-λ supernatant to primary bat nasal epithelial cells and measuring induction of ISGs (IFIT1, OAS1, RTP4) using RT-qPCR, 8 hours after stimulation. Universal interferon-α2 (Cat. #11200-1, PBL Assay Science) at 1000 U/ml was used as positive control, unmodified Lenti-X culture supernatant and Ruxolitinib (1µM, Invivogen) inhibition as negative controls. For stimulation experiments with RoIFN-λ, we used RoIFN-λ1 or RoIFN-λ3 containing supernatants at amounts needed to reach IFIT1 mRNA induction that are equivalent to induction with universal interferon-α2 (1:50-1:100 RoIFN-λ1 dilution equivalent to 1000 U/ml universal interferon-α2).

### Interferon stimulation assays

Bat and human 3D organoids grown in matrigel were collected in cold Advanced DMEM/F12 by centrifugation. 500 µl of TrypLE Expressed was added to the organoid pellet, followed by incubation for 2 minutes at room temperature. This was sufficient to obtain large organoid fragments. Equivalent amounts of Advanced DMEM/F12 were then added and organoid fragments collected by quick centrifugation. The pellet containing organoid fragments was resuspended in small intestinal or complete distal lung organoid expansion media + 10 µM Y-27632 (bat alveolar organoids without MAPK inhibitor BIRB-796). One 50 µl matrigel droplet (1-well of a 24-well plate) of a 7–10-day old organoid culture was used per 6-wells of a 96-well plate and resupended at 100 µl of medium per conditions (600 µl of medium per matrigel droplet containing organoids).

The organoid fragment suspension was then seeded at into a matrigel coated (1:50 dilution of matrigel in cold Advanced DMEM/F12 for 1-hour) 96-well plate (100 µl of organoid fragment suspension per well equals roughly 50.000 per well). Alternatively, expanding bat nasal organoids were dissociated with TrypLE and seeded as single cell monolayers onto PureCol treated wells of a 96-well plate in PneumaCult-Ex plus expansion medium at 15.000 cells per 1-well of a 96-well plates. After two to three days post seeding, diluted amounts of recombinant bat RoIFN-λ, or 100U of universal interferon-α2 or 100 ng/ml of recombinant human IFN-λ1 (Peprotech) were added for 8 hours unless otherwise indicated. RNA was harvested by removal of culture medium and addition of 200 µl of 1 M DTT (Roche) supplement Kingfisher RNA lysis buffer (Thermofisher). Lysates were stored at –70 °C for up to two weeks prior to RNA extraction. For stimulation of human organoids grown at the air-liquid interface, recombinant interferon was diluted in Gibco OptiPro serum-free medium (100ng/ml hIFN-λ1 or 1000U/ml uIFN-α2) and added to the apical chamber of the culture insert for the indicated amount of time (8 hours unless otherwise indicated). RNA was harvested by removal of culture medium from the apical and basal chamber, followed by addition of 400 µl of DTT supplement Kingfisher RNA lysis buffer and subsequently transfered to collection tubes for storage at –70 °C.

### Viruses and virus infections

All experiments involving MERS-CoV, SARS-CoV-1 and MARV infections were done in compliance with the Swedish public health agency guidelines (Folkhälsomyndigheten, Stockholm) in the appropriate biosafety level 3 (MERS and SARS-CoV-1) and biosafety level 4 (MARV) laboratories. All experiments involving Sendai virus (murine respirovirus) and VSV-EGFP infections were performed in a biosafety level 2 laboratory in compliance with the health and biosafety committee at the Vienna BioCenter. Virus strains used were Marburg virus (MARV) Musoke strain (GenBank accession number DQ217792), MERS-CoV (EMC/2012), SARS-CoV-1 (Frankfurt-1), Sendai Virus (BEI Resources NR-3227) and VSV-EGFP (Indiana strain modified to express EGFP).

For infection of distal lung and small intestinal organoids, organoid fragments were seeded on matrigel coated 96-well plates as described above for interferon stimulations. Intestinal organoids were seeded in expansion medium until confluency was reached, followed by differentiation medium for 3-4 days. For infection, the medium was removed and replaced with absorption medium (Gibco OptiPro Serum free medium, 1:100 Glutamax + 10 mM HEPES Gibco) containing virus at 50.000 PFU per-well (estimated MOI 0.5-1). After 5 hours of absorption (1 hour for Sendai virus and VSV-EGFP), the inoculum was removed and replaced with fresh organoid growth medium (that is differentiation medium for small intestinal organoids or alveolar growth medium without BIRB-796 MAPK inhibitor for distal lung organoids). RNA was harvested at indicated time-points by removing the supernatant and addition of Trizol. Trizol lysates were transferred into separate tubes and handled according to the respective biosafety protocols before storage at –70°C and RNA extraction.

For 24-well transwell grown air-liquid interface (ALI) cultures, the apical side was first washed once with PBS to remove excess mucus and dead cells. Infection was initiated by adding 100 µl of virus inoculum containing 100.000 PFU (MOI ∼0.5-1) in virus absorption medium (Gibco OptiPro Serum free medium + 1:100 Glutamax + 10 mM HEPES) to the apical chamber of ALI cultures. After incubation for 5 hours in a tissue culture incubator, the inoculum was removed, and fresh ALI medium was added to the basal chamber. At the indicated time points following infection, media was removed, and RNA was harvested by addition of Trizol and transfered to separate collection tubes for storage at –70°C and RNA extraction.

### Interferon virus protection assays

Expanding nasal, tracheal, and small intestinal organoids cultured in expansion medium (described above) were collected in cold Advanced DMEM/F12. The intact organoids where then resuspended in medium containing recombinant interferons (1000 U/ml universal IFN-α2 and functional equivalent amounts of bat interferon-λ) and incubated for 12 hours before adding different amounts of VSV-EGFP (10.000 PFUs for small intestinal organoids, estimated MOI ∼1-2; 5000 PFUs for airway organoids, estimated MOI ∼1-2). After 16 to 24 hours post infection, epifluorescent images were taken to measure virus infection (through virally encoded EGFP). In a separate experiment, expanding nasal organoids were dissociated with TrypLE and seeded as single cell monolayers in PneumaCult-Ex plus expansion medium (15.000 cells per 1-well of a 96-well plates). The next day, interferon dilutions (100U universal IFN-α2 and functional equivalent amounts of bat interferon-λ) were added for 12 hours, after which the medium was changed to PneumaCult-Ex plus expansion medium without interferons. Cells were then either immediatly infected with VSV-EGFP at an MOI of 1, or after 12 hours following medium addition (See schematics in Supplemental Figure S13C,D). After 12 or 36 hours post infection, epifluorescent images were taken to measure virus infection. EGFP image intensities were quantified with ImageJ. In brief, a grid was drawn over each image, resulting in 130 evenly sized image segments. The fluorescent intensities of each segment were then measured to represent the image intensity of each sample.

### CRISPR-Cas9 gene editing in bat small intestinal organoids

Guide RNAs targeting the Egyptian fruit bat *IRF9* gene were designed with Benchling using the NCBI IRF9 mRNA reference sequence as input. Oligos containing gRNA spacer sequences were ordered with overhangs and cloned into the lentiCRISPR v2 (Plasmid #52961) using golden gate cloning. In brief, top and bottom strand oligos (1 µl of 100 µM each) were phosphorylated with T4 PNK (New England Biolabs) for 30 minutes at 37 °C, denatured for 5 minutes at 95 °C and slowly cooled to 25 °C (0.1 °C/seconds ramp rate) for annealing. One microliter of 1:100 diluted product was used for golden gate cloning into lentiCRISPR v2-Puro (Addgene plasmid #52961) using 11 cycles of 5 minutes T4 ligase ligation at 16 °C and 5 minutes of BsmbI digestion at 37 °C, followed by a final digestion of 15 minutes at 37 °C. Two microliters of the reaction was transformed into Stbl3 E.Coli cells and grown at 34 °C. After plasmid isolation, Lentivirus was produced by transfecting 95 % confluent Lenti-X cells (Takara) grown in wells of 6-well plates with 1000 ng psPAX2 (Addgene plasmid #12260) packaging plasmid, 500 ng of pCMV-VSV-G (Addgene plasmid #8454) and 1000 ng of cloned LentiCRISPRv2-Puro plasmid using Lipofectamine (5µl P3000, 7µl L3000). After 6 hours, the media was changed to virus production medium containing Opti-MEM I (Gibco) + 5 % FBS + 1:100 Glutamax + 1:100 MEM-NEAA (Gibco) + 1:100 Sodium Pyruvate + 1:100 Penicillin/Streptomycin solution. Virus was harvested after 24 and 48 hours post transfection, filtered, and stored at –70 °C. For editing, small intestinal organoid fragments were prepared as described above for passaging, resuspended in 50 µl of undiluted lentivirus supernatant + 50 µl expansion medium + 5 µM Cyclosporin-A (Sigma) + 5 µg/ml Polybrene (Sigma) and spun at 800 rcf for 1.5 hours at 32 °C. The supernatant was removed, and organoid fragments were resuspended in 50 µl of cold matrigel and seeded as droplets into 24-well plates. Expansion medium containing + 5 µM Cyclosporin-A + 10 µM Y-27632 was added for 24 hours, then replaced with standard expansion medium. Puromycin selection (1 µg/ml) was started 60 hours post transfection. Organoids were passaged two more times in Puromycin selection media before they were used in experiments. IRF9 loss of function was determined by ISG induction using RT-qPCR in response to interferon stimulation.

### RNA extraction

For RNA extractions involving non-infected material, a semi-automated Kingfisher (Thermofisher) magnetic bead assisted protocol was used, including DNAseI (RNase free, New England Biolabs) digestion to remove genomic DNA. For virus inactivated samples in Trizol, a magnetic bead-assisted semi-automated purification workflow was performed using Direct-Zol-96 RNA kits (Zymoresearch), including DNAse I digestion to remove genomic DNA. RNA was eluted in 50 µl nuclease free water and stored at –70 °C for further use.

### RT-qPCR and analysis

Reverse-transcription on total RNA (100 to 500 ng) was performed with random hexamer/oligo-dT-Primer containing LunaScript RT Supermix (New England Biolabs) following the manufactureŕs instructions. First-strand cDNA was diluted 1:6 in nuclease free water. Two microliters were subsequently used in real-time quantitative PCR reactions containing SYBR-green based Luna universal dye qPCR mix (New England Biolabs) and gene specific PCR primers, following the ‘fast-cycling’ protocol outlined in the manufactureŕs instructions. RT-qPCR Primers were designed using IDT PrimeQuest tool, using the NCBI transcript ID of interest and “qPCR Primer and intercalating dye” as input options. For data analysis, deltaCt (dCt) values were first calculated by subtracting the Ct-value measured for the reference gene EEF1A1 from the Ct-value of a given gene measured from the same sample cDNA (e.g dCt for ACE2 in sample 1 is defined by calculating Ct-ACE2 (sample 1) minus Ct-EEF1A1 (sample 1)). Normalized relative expression values are then calculated by raising the negative dCt value of each sample by the power of 2 (2^-dCt). This represents a relative expression value of a given gene in a given sample in non-logarithmic space. Normalized relative expression values were used for visualization and statistical anaylsis in *Graphpad* prism. Primer pairs can be found in Supplemental Table 5.

### Immunofluorescence staining

Organoids were first fixed in 4 % Paraformaldehyde (Sigma) in PBS for 45 minutes to two hours at room temperature followed by three washes with PBS and storage in PBS at 2-8 °C. Organs were fixed by placing 3-5 mm small tissue pieces in 4 % Paraformaldehyde solution in PBS for 1 hour at room temperature, before transferring them to 2-8 °C for overnight fixation. PFA was removed and tissue washed three times in PBS before processing. Next, fixed samples were embedded in paraffin and sectioned at 2 µm thickness. Sample sections were dewaxed before a 30-minute citrate buffer (10mM Na-Citrate pH 6, 0.05 % Tween-20) antigen removal step in a water simmer device. After allowing for sections to cool to room temperature, samples were blocked in PBS Buffer containing 5 % Donkey Serum (Sigma) and 0.25 % Triton-X-100 (Sigma) for 30 minutes. Primary antibody dilutions in PBS containing 3 % Donkey Serum and 0.05 % Triton-X-100 were then added to samples overnight at 2-8 °C in a humidified chamber. The following day, sections were washed three times in PBS, before addition of secondary antibodies diluted in PBS containing 3 % Donkey Serum and 0.1 % Triton-X-100 or 2 hours at room temperature in a humidified dark chamber. Sections were washed three times with PBS, DNA counterstained with DAPI or Hoechst, and specimen mounted in mounting media. The following antibodies were used: anti-KRT5 (Rabbit, Sigma SAB4501651, 1:200), anti-acetylated α Tubulin (6-11B-1, SCBT, 1:500), anti-AVIL (Rabbit, Thermofisher # PA5-90703, 1:200), anti-SFTPC (Rabbit, Thermofisher #PA5-71680), anti-E-Cadherin (Mouse, BD #610182), anti-ACE2 (Rabbit SN0754, Thermofisher # MA5-32307), anti-IFN-epsilon (Mouse monoclonal, RnD Systems #MAB9147-100).

### Droplet based single-cell RNA-sequencing

Single-cell suspensions of organoids were obtained by dissociating organoids or air-liquid interface membranes containing cells in TrypLE + DNase I for 15-30 minutes at 37°C using manual pipetting in 5 minute intervals. Cells were collected by centrifugation and viable cells enriched by an Annexin-V magnetic bead dead-cell removal step following the manufactureŕs instructions. Recovered viable cells were pelleted by centrifugation, resuspended in Advanced DMEM/F12 and counted. Single-cell suspension from identical organoid formats, but different sample individuals were pooled and submitted to an in-house facility for generating 3’ gene expression libraries using the 10x Chromium Single Cell 3’ Reagent Kit v3. Sample pooling was performed to reduce batch variability and cost. Cells from individual samples could be bioinformatically separated using species (human versus bat) or single-nucleotide polymorphisms (SNP) information. Illumina 10x Libraries were sequenced on a NovaSeq S4 flowcell at 0.25 fraction of a lane per 10x library.

Sequenced reads from single cell sequencing libraries were further processed using *Cellranger* (version 7.1). First, *cellranger mkref* was used to produce a custom genome index of the *Rousettus aegyptiacus* (Assembly mRouAeg1.p) or human (Assembly GRCh38.90) reference genome. For samples with human and bat cells, a combined custom index was generated. A count matrix was then generated using *cellranger count*. *Cellbender*^112^, a recently published pipeline to remove noise in single-cell gene expression datasets, was then used to create the final filtered count matrices and barcode files. In brief, *cellbender remove-background* was run with the *raw_feature_bc_matrix.h5* file as input and the following options (--expected-cells 20000 –-total-droplets-included 30000 –-fpr 0.01 –-epochs 150). The *souporcell* pipeline^113^ (souporcell_pipeline.py) was further used to derive SNPs and assign individual cells (from *barcodes.tsv* file) to samples from aligned reads (possorted_genome_bam.bam) using the following options (-k X, whereas X is the number of expected samples; –t 8 –-skip_remap True –-ignore True). The output files, including *cellbender* filtered count matrix, barcode files, *souporcell* cluster information and *cellranger* count gem call assignment file were then used for downstream analysis in R using *Seuratv4* (4.2.1)^50, 51^. Individual *Seurat* files for bat and human cells were derived by first creating a common *Seurat* object per 10x library, addition of cell metadata from *cellranger* and *souporcell*. *Subset* was then run to obtain individual sample *Seurat* objects by using the metadata information. Multiples or unassigned cells were discarded from downstream analyses.

Samples from bat airway or small intestine organoids were integrated using *SeuratCCA*^50, 51^ (mostly adhering to the online vignette with modifications described below). First, a list of Seurat objects containing individual samples was generated. Using a loop to iterate through each object of the list, samples were then filtered to retain cells with more than 500 expressed features (genes) per cell and log-normalized (*Seurat* default option). Cell-cycle scores for each sample were calculated using G2M-/or S-Phase markers (*Seurat* vignette) and used to regress out as unwanted source of variation during gene expressing scaling performed downstream. Lastly, the top 2000 most variable features per sample were calculated. Anchors for integration were computed using *FindIntegrationAnchors* on the list containing sample objects. Data integration was performed with anchor information with *IntegrateData* and stored as an integrated Seurat object. Expression values of the integrated Seurat object were then scaled by *ScaleData* (regress out S-Phase and G2M-Phase scores information described above). PCA was performed (RunPCA), and *ElbowPlot* used to define the number of PCAs for UMAP dimensionality reduction. *FindNeighbours* and *FindClusters* with different resolution parameters (0.5-1) were used to find cell clusters. The function *FindAllMarkers* (min.pct = 0.25, log.fc=0.5, only.pos = T) was run to get a list of enriched genes for each cell cluster. Finally, we used previously defined marker genes of cell types to manually annotate cell clusters and set identities. Refined rare cell clusters were manually set by defining cells with expression cutoffs. GP2+Microfold: Ciliated/Microfold cluster cells with GP2 expression > 0; Ionocytes: Suprabasal/Ionocytes cluster cells with ASCL3 and PDE1C expression > 0; Airway brush cells: Brush and Brush/AT2 cluster cells with AVIL and POU2F3 expression > 0; Airway neuroendocrine: Suprabasal/Neuroendocrine cluster cells with CHGA and CHGB expression > 0. For differential gene expression analysis and plotting of merged datasets, raw counts were normalized with SCTransform. Analysis of human nasal and bronchial single-cell data was performed on individual datasets following the *Seurat* guided clustering vignette online. Cell clusters were identified based on known cell type specific marker genes.

For human and bat small intestinal organoids and human small intestinal tissue single-cell RNA sequencing data integration, a list of 1:1 orthologs was first determined between annotated human and Egyptian fruit bat genes. New *Seurat* object with orthologs were generated and integrated using *SeuratCCA* as described above for bat cells. External small intestinal 10x tissue single cell sequencing data was downloaded from NCBI GEO (GSE185224^68^). Ileum, duodenum and jejunum of three human donors each were included.

### Bulk RNA sequencing

Libraries for 3’end RNA sequencing were generated following the PLATE-Seq protocol with modifications^114^. Varying amounts of total RNA were mixed with 1 µl of 10 µM UMI-containing-barcoded anchored oligo-dT primers (5’Adaptor-BC(8)-UMI(12)-dT(35)VN-3’) Primer, 250 nM dNTPs mix and heated at 72°C for 3 minutes. A reverse transcription mix containing 1x First-strand buffer, 10 mM DTT, 0.15 µl Murine RNase Inhibitor (New England Biolabs) and 0.2 µl Superscript III enzyme (Thermofisher) was added to get a total volume of 20 µl. For 48-samples, twenty nanograms of RNA were used per reverse transcription reaction and scaled accordingly. Reverse transcription was performed at 50 °C for 30 minutes, followed by heat inactivation at 85°C for 10 minutes. Ten microliters of barcoded RT reaction were then pooled into a single tube. 1 µl per 50 µl pool volume of Thermolabile Exonuclease I (New England Biolabs) was then added and incubated for 4 minutes at 37 °C, 1 minute at 80 °C. The cDNA from the pool was purified using in-house AmpureXP-like beads at 1:1.4 sample to bead ratio and eluted in 50 µl nuclease free water. Ten microliters each of 1 M NaOH and 0.5 M EDTA pH 8 were added to the purified sample and heated at 65 °C for 15 minutes to hydrolyze RNA in the RNA:cDNA hybrids. The reaction was purified using a ZymoResearch Oligo Clean & Concentrator Kit following the manual instructions for cDNA clean-up and eluted in 17 µl. A 23 µl reaction containing purified first-strand cDNA, 2.5 µl of New England Biolabs Buffer 2, 1 µl of 100 µM adaptor-random-hexamer primer, 200 nM dNTPs mix was heated to 95°C for 1 minute, followed by slow cooling to room temperature (0.1°C/second) in a PCR thermocycler. Second-strand synthesis was performed by addition of 2 µl Klenow-Fragment DNA Polymerase (New England Biolabs) and incubation for 15 minutes at 30°C. The reaction products were purified with a Zymo Research DNA Clean & Concentrator-5 kit and eluted in 20 µl nuclease free water. A final limited cycling PCR was performed using dual-indexing Illumina Primers (5 µM each primer), Q5 NEBNext® Ultra™ II Q5® Master Mix (New England Biolabs) for 10-14 cycles. The reaction was gel-purified, excising fragments between 200-900bp and collected in 20 µl elution buffer (10mM TrisCl pH 8). Libraries were sequenced on an Illumina NovaSeq S4 lane using 150bp pair-end mode.

Fastq read files were processed using the BRB-seqTools pipeline^115^ (https://github.com/DeplanckeLab/BRB-seqTools). First, read-1 was trimmed to 25 nucleotides, while read-2 was aligned to the *Rousettus aegyptiacus* (Assembly mRouAeg1.p) or human (Assembly GRCh38.90) reference genomes using STAR. *BRBSeq-CreateDGE* Matrix was then used to create an UMI-count sample file matrix from the aligned read-2 bam file and trimmed read-1 fastq file containing UMI and sample barcode information. The gene expression count file was used for downstream *edgeR*^116^ differential gene expression analysis in R. Lowly or non-expressed genes were removed from subsequent analysis (that is genes with less than 2 counts per million (CPM) in less than 50 % of replicates of a given sample group (e.g mock treated, interferon treated). TMM normalization was used to assess normalized log2-counts per million (CPM). A *genewise negative binomial generalized linear model* model was subsequently fit to the count data providing sample group information using glmQLFit (*edgeR* vignette). Differentially expressed genes (DEG) were determined (using the model fit above and edgeR *glmQLFTest*) and filtered for significantly up-/or downregulated genes (PValue < 0.05). Differential gene expression analysis tables were used for visualization and statistical analysis in *Graphpad Prism* (version 9). Gene ontology enrichment analysis of upregulated genes was performed though clusterProfiler^117^ in R. Exclusive and overlapping sets of differentially expressed genes were visualized with *eulerr* (eulerr.co).

### Re-analyses of *in vivo* MARV Egyptian fruit bat infection data

Gene expression re-analysis from MARV infected Egyptian fruit bats was performed by downloading the published nCounter normalized data from Guito et al.^7^ A data subset was generated containing expression of interferon stimulated genes^118^ and Marburg virus genes for selected tissue samples, including skin at the inoculation site, liver, spleen and colon. A normalized expression fold change was defined by dividing the maximum observed value for each gene by the minimum, followed by log2 transformation. Highly dynamic genes were then selected by ranking them based on the normalized expression fold change and taking the top 25% of the resulting list for downstream analyses. This list was further sub-selected by only including genes with a maximum normalized expression value falling within the top 25% most highly expressed genes across all sample for a given tissue (this removes genes with a large dynamic range that are lowly expressed in general and therefore not of interest to the analysis). The final list contains genes with moderate to high maximum expression and large dynamic expression range throughout the infection time course.

### Data and code availability

Raw and processed sequencing data, as well as codes to generate processed data and visualizations will be openly shared upon publication and will be available upon request.

### Reagent availability

All reagents including organoids can be shared upon request with a material transfer agreement.

