## Supplementary figures and images for "Reconstructing bat antiviral immunity using epithelial organoids"

### Supplementary Movie 1

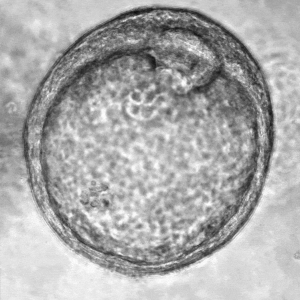

### Supplementary Movie 2

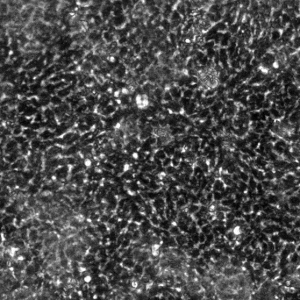
